# A Genetic Mechanism Linking Hippo Signaling to Dorsoventral Patterning for Control of Head and Eye Development

**DOI:** 10.64898/2026.01.24.701508

**Authors:** Basavanahalli Nunjundaiah Rohith, Neha Gogia, Arushi Rai, Amit Singh, Madhuri Kango-Singh

## Abstract

The integration of growth and patterning in developing tissues is a complex process involving both intrinsic and extrinsic cues. The Hippo pathway, a conserved regulator of organ size, controls growth and patterning in *Drosophila*, including the development of the eye-antennal imaginal disc into adult structures. Defective Proventriculus (Dve), a SATB1/2 ortholog and K-50 type transcription factor regulates dorsal-ventral (DV) patterning during *Drosophila* eye development. Dve works with Wingless (Wg) to suppress eye development and promote head cuticle fate, thereby influencing the positioning of eyes and interocular distance. Our study investigates the role of the Hippo effector Yorkie (Yki), and *dve* in coordinating growth and patterning during eye development, specifically focusing on the regulation of the head cuticle domain. Here we show that Hippo signaling, mediated by Yki, regulates the size of the head cuticle domain and morphogenetic furrow (MF) progression, and that Dve suppresses Yki activity in the dorsal head region. Furthermore, Yki regulates several DV patterning genes like *pnr*, *wg* and *mirr*, to coordinate eye and head development. Mutations in mammalian orthologs of *yki*, *dve*, *pnr* and *wg* are associated with facial dysmorphia and several developmental disorders. Our studies thus reveal new genetic mechanisms by which growth and patterning are coordinated for head and eye development across species.

## Introduction

The integration of growth with patterning is a complex developmental process in which cells within a developing field respond to intrinsic and extrinsic cues to regulate cell division, specification and differentiation to generate tissues and organs in a stereotypical manner (Guo et al., 2025; Singh et al., 2012). Growth is a fundamental process in all metazoans and is governed by intricate regulatory mechanisms that control cell proliferation and survival (Guo et al., 2025; Kango-Singh and Singh, 2009). These processes are highly conserved across organisms due to the conservation of core developmental patterning and signaling pathways (Housden and Perrimon, 2014; Natarajan et al., 2006). *Drosophila melanogaster* is a well-studied model for studying intricate developmental processes due to its conserved pathways and powerful genetic tools (Bier, 2005; Verghese et al., 2020).

The *Drosophila* eye-antennal imaginal disc develops into the adult eye, head and antenna; and serves as a robust model to study developmental patterning (Haynie and Bryant, 1986; Kumar, 2023; Singh et al., 2012; Weasner and Kumar, 2022b). Eye antennal progenitors arise from the embryonic head ectoderm and develop into the eye-antennal epithelial imaginal discs that give rise to the compound eyes, antennae, ocelli, the head cuticle and palpus of the adult fly (Haynie and Bryant, 1986; Singh et al., 2012; Weasner and Kumar, 2022a). The growth and differentiation mechanisms involved in the transition of the monolayered eye-antennal imaginal discs to the adult structures are tightly controlled by interactions among morphogens, signaling pathways and transcriptional regulators (Bessa et al., 2002; Chang et al., 2001; Lopes and Casares, 2010; Ma and Moses, 1995; Singh and Choi, 2003; Singh et al., 2012). Although much is known about the mechanisms that regulate the differentiation of photoreceptor neurons in the wake of the morphogenetic furrow (MF) that sweeps across the eye disc from posterior to anterior (Ma and Moses, 1995; Treisman and Rubin, 1995; Weasner and Kumar, 2022a), the regulation of other regions of the eye discs including the head cuticle domain remains poorly understood.

The Hippo pathway is an evolutionary conserved signaling pathway for regulation of patterning and growth to control organ size in both *Drosophila* and mammals (Guo et al., 2025; Kango-Singh and Singh, 2009). The upstream regulators of the pathway impinge on a core-kinase cascade comprised of the serine-threonine kinase Hippo (Hpo, ortholog of mammalian Sterile-20 like kinases 1 & 2, MST1/2) which partners with the adaptor protein Salvador (Sav, hSAV1) to activate the downstream serine-threonine kinase Warts (WTS) *a.k.a.* Large Tumor Suppressor (LATS, hLATS1/2). LATS in turn, partners with the adaptor protein Mob as Tumor Suppressor (MATS, hMOB1A/B) that potentiates its catalytic activity. This complex phosphorylates and inactivates the transcriptional co-activator Yorkie (Yki), ortholog of human Yes-Associated Protein ½ (YAP1/2) and Transcriptional coactivator with a PDZ-binding domain (TAZ). Activated Yki/YAP partners with TEA/ATTS family of transcriptions factors like Scalloped (Sd, TEAD1/2/3/4) or MEIS-family transcription factor Homothorax (Hth, hMEIS1) or zinc-finger transcription factor Tea-shirt (Tsh, hTSH3). The Hippo pathway is known to regulate overall organ size, and changes in pathway activity perturb developmentally regulated organ size (Guo et al., 2025; Kango-Singh and Singh, 2009). Over-expression of Yki results in its nuclear translocation and association with the TEAD-family transcription factor Scalloped (Sd) or Meis-family transcription factor Homothorax (Hth) (Peng et al., 2009; Slattery et al., 2013; Zhang et al., 2011). These transcriptional complexes activate target genes that promote cell division and suppress cell death. Recently, multiple roles for the Hippo pathway in developmental patterning have been described in *Drosophila* and higher vertebrates (Barry and Camargo, 2013; Dong et al., 2007; Gangwani et al., 2020; Zhao et al., 2008).

During eye development, dorsal-ventral (DV) patterning plays a key role in defining the first lineage restriction by the expression of Pannier (Pnr) in the dorsal eye margin (Maurel-Zaffran and Treisman, 2000; Oros et al., 2010; Pichaud and Casares, 2000; Singh et al., 2006; Singh et al., 2011). We recently identified *defective proventriculus* (*dve*) as a Pnr-regulated dorsal fate selector gene (Puli et al., 2024). Dve is a K50 homeodomain-containing transcription factor in *Drosophila* that is highly conserved from insects to humans, and homologous to the human Special AT-rich sequence-binding proteins (SATB1/2) (Pavan Kumar et al., 2006; Singh et al., 2012). SATB1 regulates chromatin to provide docking sites for transcription factor and chromatin modifying enzymes, thereby controlling gene expression (Pavan Kumar et al., 2006). Initially, *dve* was shown to be required for embryonic gut development (Fuß and Hoch, 1998; Nakagoshi et al., 1998). Subsequent studies showed that *dve* also regulates development of other organs including legs (Shirai et al., 2007), wings (Nakagoshi et al., 2002; Sugimori et al., 2016), and eyes (Puli et al., 2024). In the eye discs in *Drosophila*(and other insect species), Dve is expressed in the dorsal head vertex region, where it regulates the expression of *wingless* (*wg*), a secreted morphogen expressed along the anterolateral eye margins, and is involved in determining the eye versus head fate (Peng et al., 2017; Puli et al., 2024). Dve and Wg suppress eye fate by downregulating retinal determination genes and promoting head cuticle development. Loss of function of *dve* results in dorsal eye enlargement and gain of function of *dve* causes eye suppression as well as reduced size of the head capsule in flies (Puli et al., 2024). Overall, Dve plays a key role in DV patterning by controlling the positioning of eyes on the head and defining interocular distance during development (Puli et al., 2024).

However, how growth and patterning mechanisms are coordinated during development is not fully characterized. In particular, the regulation of the head cuticle domain that determines the spacing between the two eyes remains largely unexplored. Here, we investigated a putative role of Hippo signaling and its effector *yki* and *dve* in determining the optimum growth of eye and head cuticle domains for normal tissue homeostasis. We show that Hippo signaling affects size of the head cuticle domain size and MF progression during eye development. We further show that Dve- a dorsal fate selector- suppresses Yki activity in the dorsal head region. Further, we found that *dve* itself is not transcriptionally regulated by Yki in the eye disc, however, *pnr* the upstream regulator of Dve and *mirror* (*mirr*), a gene from the Iroquois-complex (Iro-C) that acts downstream of Wg in the dorsal eye, as new transcriptional targets of Yki. Taken together, our studies support a model in which Dve controls head cuticle domain size by antagonizing Yki within the Hippo pathway, and Yki functions upstream of *pnr* and transcriptionally regulates genes involved in DV patterning and growth regulation during head cuticle expansion and retinal specification. These studies provide a direct mechanistic link between growth control and axial patterning during head-eye morphogenesis, and given the conservation genes and developmental pathways, our studies are relevant for understanding the etiology of conditions involving craniofacial malformations and dysmorphia in higher vertebrates including humans.

## Results

### Hippo signaling affects the size of the head cuticle domain and MF progression during eye development

Loss of function of *dve* in the eye (using *ey-FLP, FRT42D cl w+*) resulted in dorsal eye enlargements and suppression of Wg expression (Puli et al., 2024). Compared to wild-type eye-imaginal discs (Fig. 1A, A’) and adults (Fig. 1B), *dve-GAL4 UAS-GFP* (referred to as *dve>GFP*, Fig. 1C, C’) shows Dve expression (GFP-positive cells in Fig. 1C, grey in C’) in about 150-200 cells in the dorsal head vertex region of the eye disc. These cells do not overlap with the differentiated photoreceptors marked by antibodies against ELAV, a pan-neural marker in *Drosophila* (Fig. 1C red). Upregulation of Hippo pathway activity under *dve-GAL4* by over-expression of Hpo (*dve>GFP, Hp*o Fig. 1E, E’) or Wts (*dve>GFP, Wts* Fig.1G, G’) resulted in loss of Dve domain (Fig. 1E’, G’) and concomitant loss of antenna and head cuticle regions in the discs (Fig. 1E, G) (Puli et al., 2024). In addition, these discs showed remarkable extension of eye by formation of ectopic or elongated furrows leading to defects in patterning in the entire disc (Fig. 1E, G). These defects in differentiation and patterning can be further tracked post-metamorphosis in adults or pharates. Compared to wild-type (Fig. 1B) or *dve>GFP* adult flies (Fig. 1D), the *dve>GFP, Hpo* (Fig. 1F) or *dve>GFP, Wts* (Fig. 1H) animals fail to eclose, and pharate adults show defects in head eversion and/or head capsule formation. As a result, the differentiated eyes remain trapped within the thorax. Together, these data suggest that overexpression of Hippo pathway components in the Dve domain affects eye patterning and affects the size of the Dve expression domain. Therefore, we asked if the Hippo pathway modulates the head versus eye fate decisions in the developing eye antennal discs.

**Figure 1.**
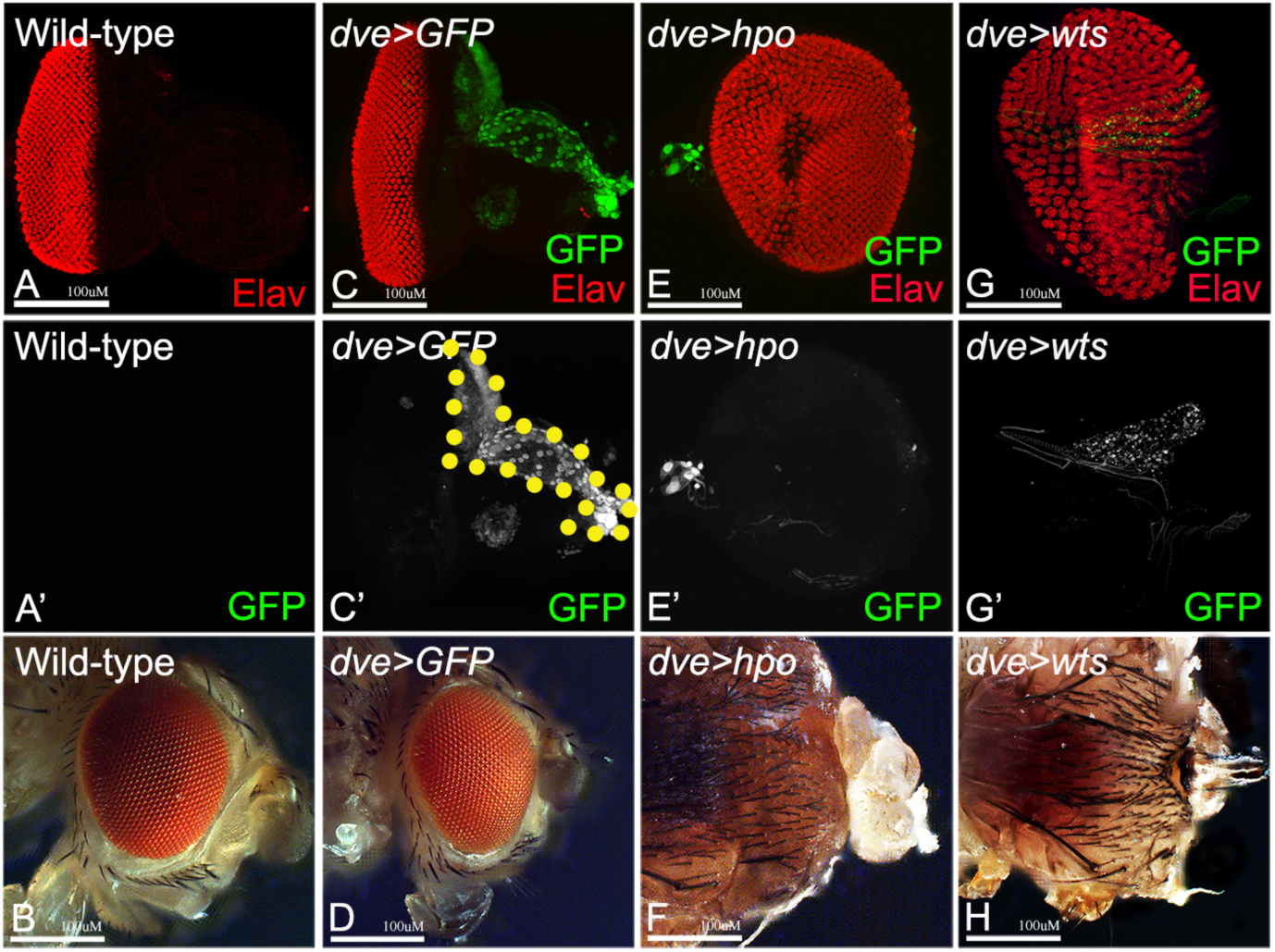
Upregulation of Hippo pathway activity affects eye and head development. Comparison of *dve* expression domain size and its effects on eye and head development is shown for (A,B) wild-type, (C,D) *dve-GAL4>UAS GFP* (GFP, green), (E,F) *dve-GAL4> UAS GFP, UAS hpo*, and (G,H) *dve-GAL4> UAS GFP, UAS wts.* Panels A, C, E, G show confocal images of third instar eye imaginal discs stained for the pan-neuronal marker ELAV (red), and *dve-GAL4* expression domain marked by *UAS GFP* (green in C,E,G; grey in C’,E’,G’). Panels (B, D, F, H) show adults flies from corresponding genotypes. In panel C’, wild type *dveGAL4* expression domain is marked by UAS GFP and is outlined by yellow-dotted line. Scale bar = 100µM

### Hippo activity affects dorsal head domain size

To test if the Dve domain is indeed affected by manipulating Hippo pathway function, we tested the effects of depleting Hippo pathway activity (Fig. 2). To do so, we overexpressed the physiological- (*UAS-Yki*) and hyperactivated- (*UAS-Yki^3SA^*) forms of Yki that are known to mimic the loss of function effects of the Hippo pathway (Oh and Irvine, 2009). Compared to the controls (*dve>GFP*) (Fig. 2A-A”), over-expression of the physiological form of Yki in the Dve domain (*dve>GFP, Yki,* Fig. 2B-B”) resulted in a significant increase in size of the Dve expression domain (Fig. 2G), and suppression of the morphogenetic furrow leading to formation of smaller eyes (Fig. 2H). Interestingly, over-expression of hyperactivated form of Yki (*dve>GFP, Yki^3SA^*) further enhanced these defects in the discs (Fig. 2C-C”) and adults (Fig. S1A-C). Quantification of the domain size (area of GFP expression in each genotype) shows that overexpression of both forms of Yki caused significant expansion in the Dve expression domain (Fig. 2G) and decrease in eye size due to suppression of the morphogenetic furrow progression both in the dorsal and ventral margin (Elav expression in Fig. 2B-C, quantified in Fig. 2G, H). Next, we tested the effect of downregulation of Yki on the Dve expression domain (Fig. 2D-D”). Compared to wildtype control (*dve>GFP*, Fig. 2A), downregulation of Yki (*dve>GFP, Yki^RNAi^*, Fig 2D) showed no significant effect on the domain size (quantified in Fig. 2G), and mild extension of the morphogenetic furrow (MF) (Fig. 2D” marked with arrowhead, quantified in 2H). Consistent with this, loss of function clones of *hpo* (*FRT42D hpo^42-47^/FRT82B cl w+* in Fig. S1D) or *wts* (*FRT82B wts^X1^/ FRT82B cl w+* in Fig. S1E) showed expansion of the Dve domain (green, grey in Fig. S1D’, E’) and concomitant suppression of the MF at the margins of the eye disc (Fig. S1D”, E” marked with arrowheads). Thus, Dve domain size is significantly affected by gain- or loss of Hippo signaling.

**Figure 2.**
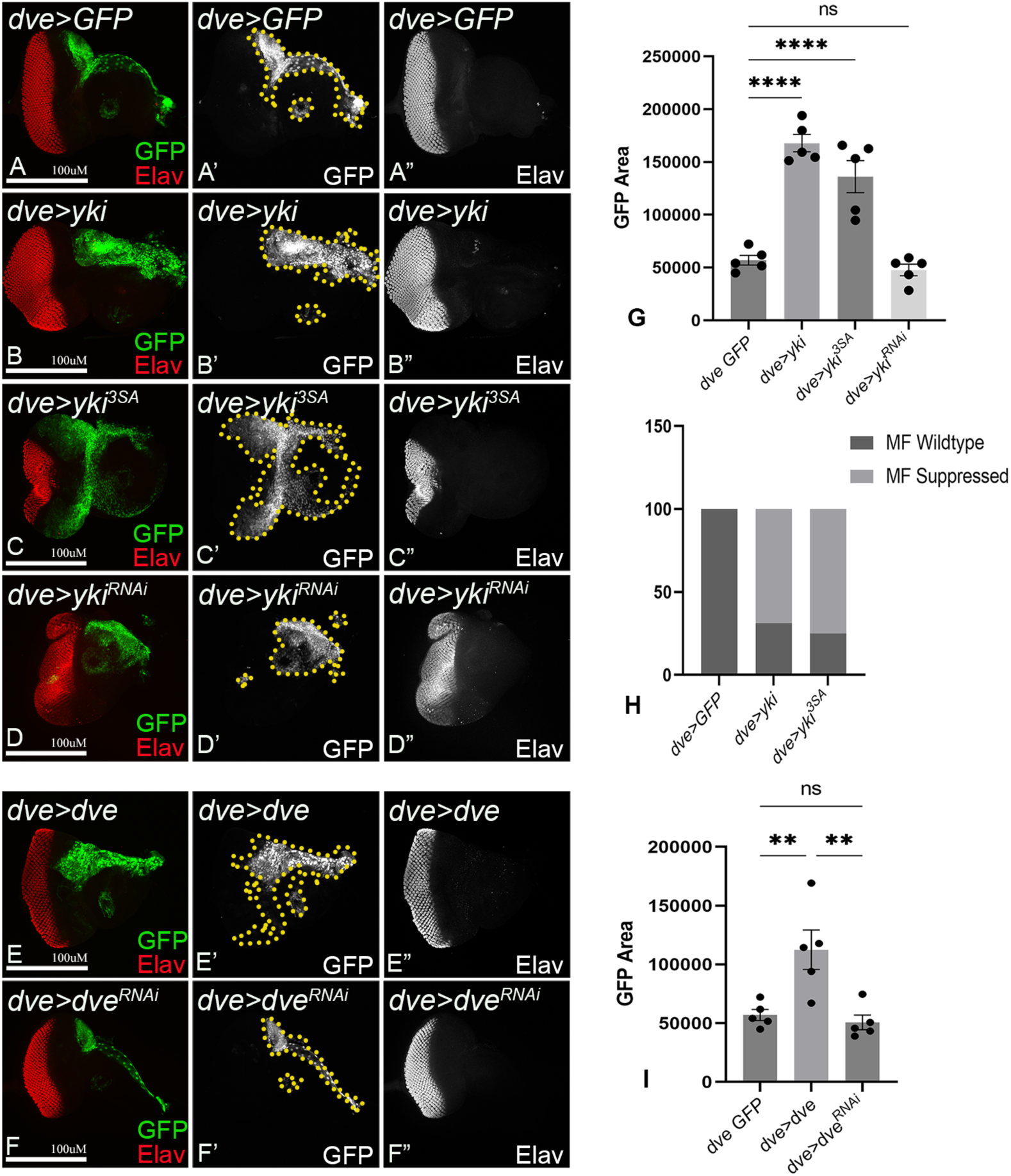
Modulating Hippo activity or Dve levels affects the domain size of the head cuticle and eye. Panels show the effects of modulating Hippo pathway activity ( A-D, quantified in G,H) and Dve expression (E-F, quantified in I) on Dve expression levels, domain size and eye size. Eye size is compared based on ELAV expression (red in A-F, grey in A”-F”) whike the size of the *dve* domain, which affects head cuticle size, was assessed by GFP expression (*dveGal4>GFP*, green in A-F, grey in A’-F’). Effects of modulating Hippo activity are shown in panels A-D and quantified in G-H. (A-D) Confocal images of third instar eye imaginal discs from (A) *dve-GAL4>UAS GFP* (*dve>GFP)* control, (B) *dve-GAL4>UAS GFP, UAS yki* (*dve>GFP, yki*), (C) *dve-GAL4>UAS GFP, UAS yki^3SA^* (*dve>GFP, yki^3SA^*), and (D) *dve-GAL4>UAS GFP, UAS yki^RNAi^* (*dve>GFP, yki^RNAi^*). ELAV (red in A-D, grey in A”-D’) expression marks differentiated photoreceptors and was used to measure eye size. The *dve* domain is outlined by GFP expression (marked by yellow dotted line in panels A’-D’). Graphs in G,H. show quantification of GFP domain size (G), and effect on overall eye size due to suppression of MF progression (H). (E-F) Panels show effects of Dve modulation. Confocal images of third instar eye imaginal discs from (E) *dve-GAL4>UAS GFP UAS dve (dve>GFP, dve)* and (F) *dve-GAL4>UAS GFP UAS dve^RNAi^*(*dve>GFP, dve^RNAi^*) showing expression of *ELAV* (red in E-F; and grey in E”-F”) that marks differentiated photoreceptors, and GFP (green in E-F, grey in E’-F’ outlined in yellow dotted line) that marks the Dve domain. Graph in I shows quantification of head domain size (GFP). Scale bar = 100µM.

Next, we checked if modulation of Yki binding transcription factor Sd also affected head cuticle domain size or MF progression in the eye disc (Fig. S1). Compared to wild-type (*dve>GFP*, Fig. S1F), overexpression of Sd (*dve>GFP, Sd*, Fig. S1 G-H) showed significant reduction in the Dve domain size (quantified in Fig. S1J) whereas RNAi mediated knockdown of Sd (*dve>GFP, Sd^RNAi^*, Fig. S1I) showed statistically insignificant differences in the Dve domain size (Fig. S1J). In addition, overexpression of Sd showed interesting expansion of the MF at the dorsal (Fig. S1G”, H”) and ventral margins (Fig. S1H”), a phenotype that is opposite to the effects of Yki overexpression where the MF was suppressed. Together, these data confirmed that the manipulation of the Yki/Sd complex affects both head cuticle domain size and MF progression in eye discs.

To assess if changes in domain size are sensitive to Dve itself, we tested if over-expression of Dve (*dve>GFP, dve*, Fig. 2E) or its downregulation (*dve>GFP, dve^RNAi^*, Fig. 2F) affects Dve domain size. We noticed an increase in the Dve domain size when we over-expressed Dve (Fig. 2E, quantified in Fig. 2I), which also resulted in reduced growth of eye imaginal discs which appeared smaller compared to wild type. The Dve overexpressing larvae (*dve>GFP, dve*) developed into adult flies with smaller heads and reduced or no eyes (Puli et al., 2024). In contrast, down regulation of Dve (*dve>GFP, dve^RNAi^*, Fig. 2F, quantified in Fig. 2I) showed no significant change in Dve domain size or morphogenetic furrow progression as compared to wild type. Together, these data suggest that the differentiation of the photoreceptor neurons and the overall dorsal head domain controlled by Dve function in the developing eye imaginal discs are very sensitive to the activity of Hippo effector Yki and the DV patterning gene Dve.

### Dve suppresses Yki activity in the dorsal head region

The Hippo pathway effector Yki transcriptionally controls several genes involved in the regulation of organ size, including *expanded* (*ex*), *death-associated inhibitor of apoptosis 1 (diap1)*, *bantam miRNA* and *hth,* which are well-characterized Yki/Sd dependent target genes (Zhang et al., 2011). First, we checked the expression of an *ex*-reporter (*ex^697^-lacZ*) (Boedigheimer and Laughon, 1993) in the head cuticle domain where Dve is expressed. Compared to control eye discs (*dve>GFP*, Fig. 3A) that show normal expression of *ex-lacZ* (Fig. 3A’), Dve overexpression (*dve>GFP, dve)* (Fig. 3B) showed reduced but not significant differences in the *ex-lacZ* reporter expression (Fig. 3B’) in the head cuticle domain (GFP in Fig. 3B, quantified in Fig. 3D). As expected, overexpression of Yki (*dve>GFP, Yki*) caused significant upregulation of *ex-lacZ* expression (Fig. 3C, quantified in D). Next, we tested if Dve exhibits any domain-specific constraints in its ability to modulate Yki activity in the developing eye. To do so, we ectopically expressed Dve using *GMR-GAL4*, which drives expression in the differentiating retinal neurons that do not normally express Dve during development (Fig. 3E-G). Compared to controls (*GMR*>, Fig. 3E), ectopic expression of Dve did not significantly affect *ex-lacZ* expression (*GMR>dve*, Fig. 3F, quantified in Fig. 3G). Taken together, these data suggest that *ex-lacZ* expression is not significantly affected by *dve* overexpression in the eye discs. Next, we tested expression of DIAP1 an important cell survival factor transcriptionally regulated by the Hippo signaling pathway [1]. Compared to control (*dve>GFP,* Fig. S2A, quantified in E), discs over-expressing *dve* (*dve>GFP, dve*) showed no significant difference in DIAP1 expression (Fig. S2B, quantified in E). Taken together, our data suggests that within the head vertex region (Dve domain), Dve does not significantly affect DIAP1 expression, whereas in the photoreceptor cells downregulation of Dve can lead to expansion/enlargement of eyes, suggesting that optimum *dve* and *yki* activity are required to make organs of proportionate size. Next, we tested Yki target genes that regulate cell death, for example, the *microRNA bantam* using the GFP-labelled *bantam sensor* (Fig. S2). The *bantam-sensor* reports inverse profile of *miRNA bantam* activity such that high GFP levels correspond to low *miRNA bantam* expression, and *vice versa* (Thompson and Cohen, 2006). Compared to wild-type eye discs (Fig. S2F), overexpression of *dve (dve>dve)* (Fig. S2G) resulted reduced eye size, however, no obvious effect on expression of the *bantam sensor* was seen (compare Fig. S2F’ with Fig. S2G’). In contrast, knockdown of *dve* (*dve>dve^RNAi^*) showed enlargement of the photoreceptor domain (Fig. S2H) as compared to the control (Fig. S2E), and a concomitant expansion of the *bantam sensor* expression (Fig. S2H’). These results show that modulation of *dve* directly affects photoreceptor differentiation and eye size but does not specifically affect *bantam sensor* expression.

**Figure 3.**
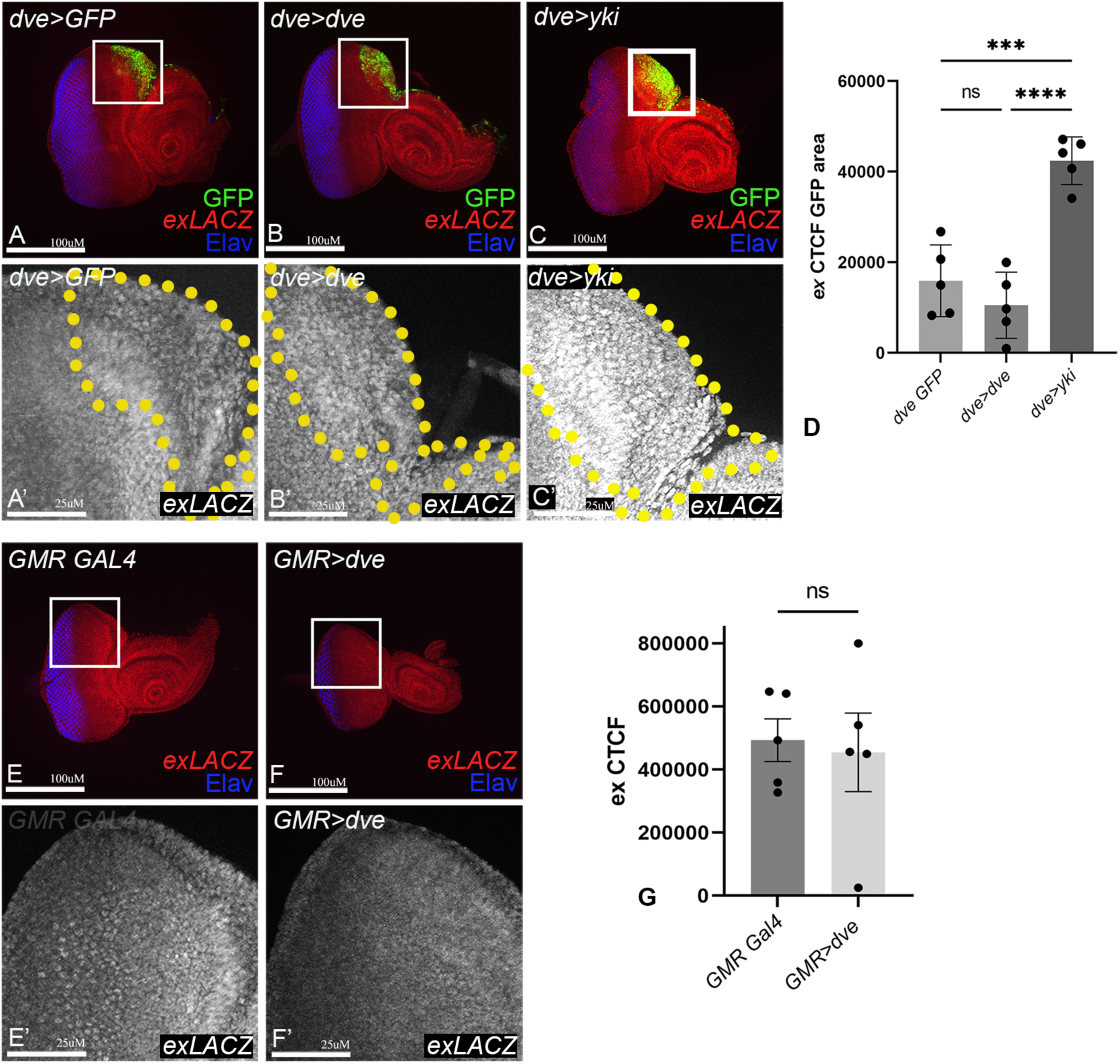
Yki target gene *expanded* is not significantly affected by Dve. (A-C) Confocal imaged of third instar eye discs showing expression of the Yki reporter gene, *ex^697^-lacZ* (red in A-E, grey in A’-E’) are shown: (A) *ex^697^-lacZ dve-GAL4 UAS GFP* control (*ex-lacZ dve>GFP*), (B) *ex^697^-lacZ dve-GAL4 UAS GFP UAS dve* (*ex-lacZ dve>GFP, dve*), (C) *ex^697^-lacZ dve-GAL4>UAS GFP UAS yki* (*ex-lacZ dve>GFP, yki*). (D) Graph shows quantification of *ex^697^-lacZ* reporter expression (CTCF). (E-G) Panels show eye discs from (E) *ex^697^-lacZ GMR-GAL4* (*ex-lacZ GMR>*) and (F) *ex^697^-lacZ GMR-GAL4>UAS GFP UAS dve* (*ex-lacZ GMR>GFP, dve*). (G) Quantification of the *ex^697^-lacZ* reporter expression (CTCF) is shown in graph in G. For A,B,C,E,F Scale bar = 100µM, for A’,B’,C’,E’,F’ Scale bar = 25µM

Hth is a transcription factor that binds with *yki* and regulates Sd-independent target genes in the eye disc (Zhang et al., 2011). Hth plays multiple roles as a negative regulator of the eye fate and a positive regulator of proliferation of eye precursor cells (Bessa et al., 2002; Singh et al., 2011; Wang and Sun, 2012; Wittkorn et al., 2015). Hth expression normally overlaps the *dve-*GAL4 domain in dorsal head vertex region (*dve>GFP*, Fig. 4A). Consistent with its role as target gene of Yki, Yki overexpression (*dve>GFP, yki)* induced Hth expression (Fig. 4B, quantified in Fig. 4E). Interestingly, Hth is suppressed in discs where Dve (Fig. 4C, quantified in Fig. 4E) or Dve and Yki are co-expressed (Fig. 4D, quantified in Fig. 4E). Taken together, these data suggests that Dve acts downstream of Yki and suppresses Yki activity in the dorsal head region.

**Figure 4.**
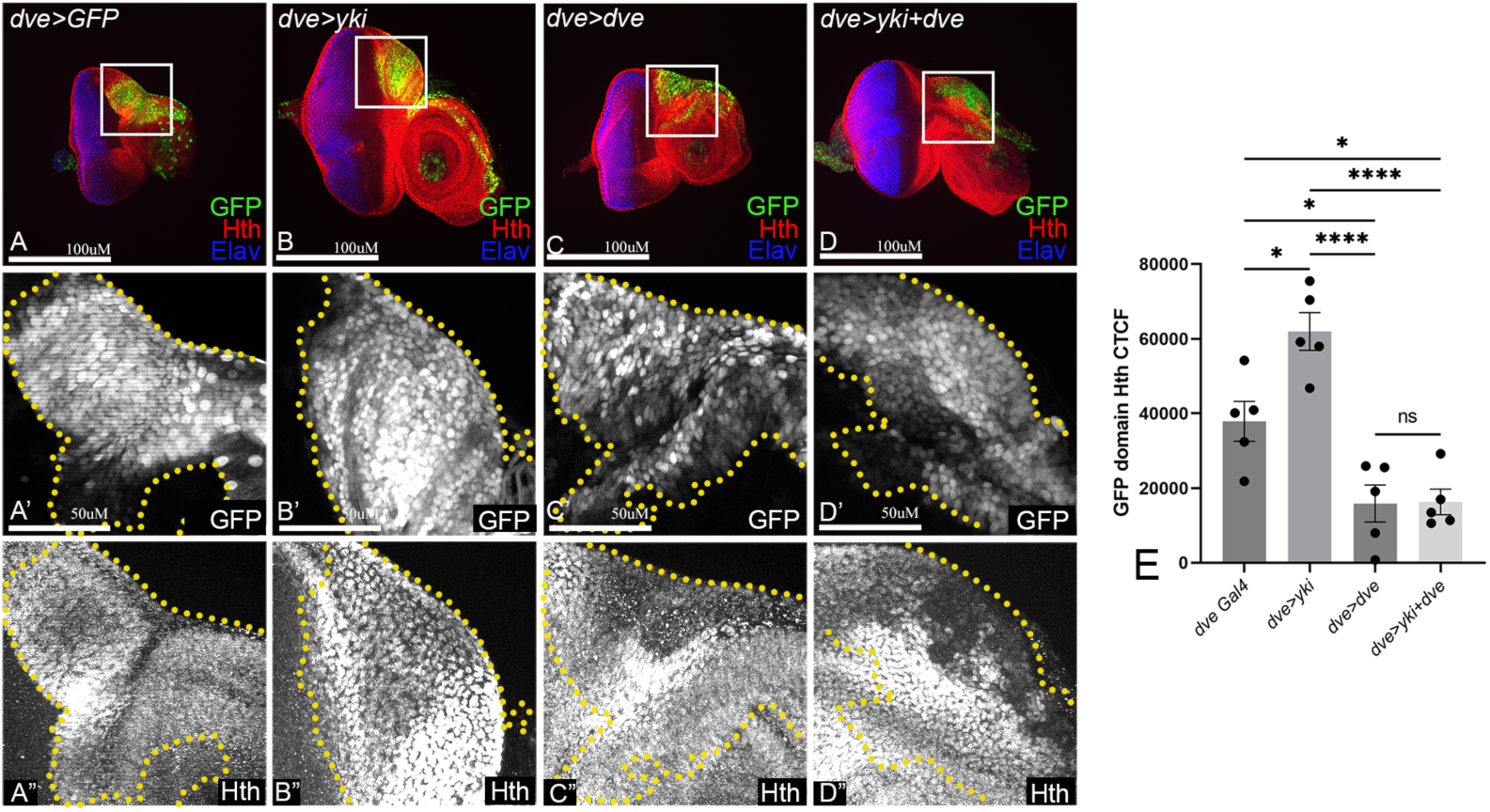
Dve suppresses Yki activity in the dorsal head region. Panels show confocal images of third instar eye discs stained for Hth (red in A-D, grey in A”-D”) for the following genotypes: (A) *dve-GAL4>UAS GFP* (*dve>GFP*) control, (B) *dve-GAL4>UAS GFP, UAS yki* (*dve>GFP, yki*), (C) *dve-GAL4>UAS GFP UAS dve* (*dve>GFP, dve*) and (D) *dve-GAL4>UAS GFP UAS dve UAS yki* (*dve>GFP, dve, Yki*). The *dve* expression domain (*dveGal4>GFP*) marks dorsal head region and is shown in green (A-D) and grey (A’-D’). (E) Quantification of Hth expression (CTCF) in the indicated genotypes is shown. For A,B,C,D Scale bar = 100µM, and for A’,B’,C’,D’ Scale bar = 50µM

### *dve* is not transcriptionally regulated by Yki in the eye disc

RNAseq studies have reported that *dve* may be a target of Yki/Sd transcriptional complexes (Zhang et al., 2017). We, therefore, tested if Yki directly regulates *dve* transcription in the eye discs. We overexpressed Yki (*UAS-yki*) and its hyperactivated form (*UAS-yki^3SA^*) in the eye using *GMR-GAL4* and *bi-GAL4* and tested effects on *dve-lacZ* expression, a reporter for *dve* transcription (Fig. S3). We mis-expressed *yki* in the differentiating photoreceptor cells using GMR-GAL4 (Fig. S3A-C). Compared to controls (Fig. S3A,B), over-expression of *yki* (*GMR>yki*, Fig. S3C) did not trigger ectopic expression of *dve-lacZ* in the GMR domain. *bi-GAL4* drives expression of UAS-tagged genes along the dorsal and ventral margins of the eye (Fig. S3D, GFP). In the *bi>GFP* controls (Fig. S3D), *dve-lacZ* expression is restricted to its normal domain in the dorsal head vertex region (Fig. S3D’). In comparison, overexpression of Yki (*bi>GFP, yki*, Fig. S3E) or its hyperactivated form (*bi>GFP, yki^3SA^*, Fig. S3F) showed the activation of Yki in dorsal or ventral margins (marked by GFP in Fig S3E’’’, F’’’) does not induce *dve-lacZ* expression (Fig. S3E’,F’). Thus, unlike the wing discs (Zhang et al., 2017), Yki may not directly regulate *dve* expression in the eye imaginal discs. Since transcriptional reporter may not fully reflect the *in-vivo* effects of Yki over-expression, we tested *dve* expression using qRT-PCR and found that in the eye discs Yki overexpression does not induce *dve* expression (Fig. S3G, H). As expected, over-expression of Dve (Fig. S3G,H) significantly induced *dve* transcript levels as compared to controls (*dveGal4>UASGFP*). Taken together, these results suggest that Yki does not transcriptionally regulate *dve* during eye imaginal disc development.

### Dve acts downstream of Yki in regulating dorsal head domain size

To further explore the interactions between Dve and Yki, we performed genetic interaction experiments. We hypothesized that Yki and Dve function epistatically to regulate shared transcriptional targets like *wg* that control growth via Hippo signaling and DV patterning. In a simple epistatic model, Yki may act upstream of Dve, or *vice-versa*. Using multiple GAL4 drivers like *dve-GAL4, bi-GAL4, GMR-GAL4* and *ey-GAL4* we investigated if temporal effects due to differences in GAL4 activation affect disc growth. Therefore, we investigated if genetic interactions between Yki and Dve using early (before differentiation) GAL4 drivers (e.g., *ey-GAL4*) show different effects on disc growth and differentiation as compared to later GAL4 drivers (e.g., *dve-GAL4* and *GMR-GAL4*) (Fig. 5).

**Figure 5.**
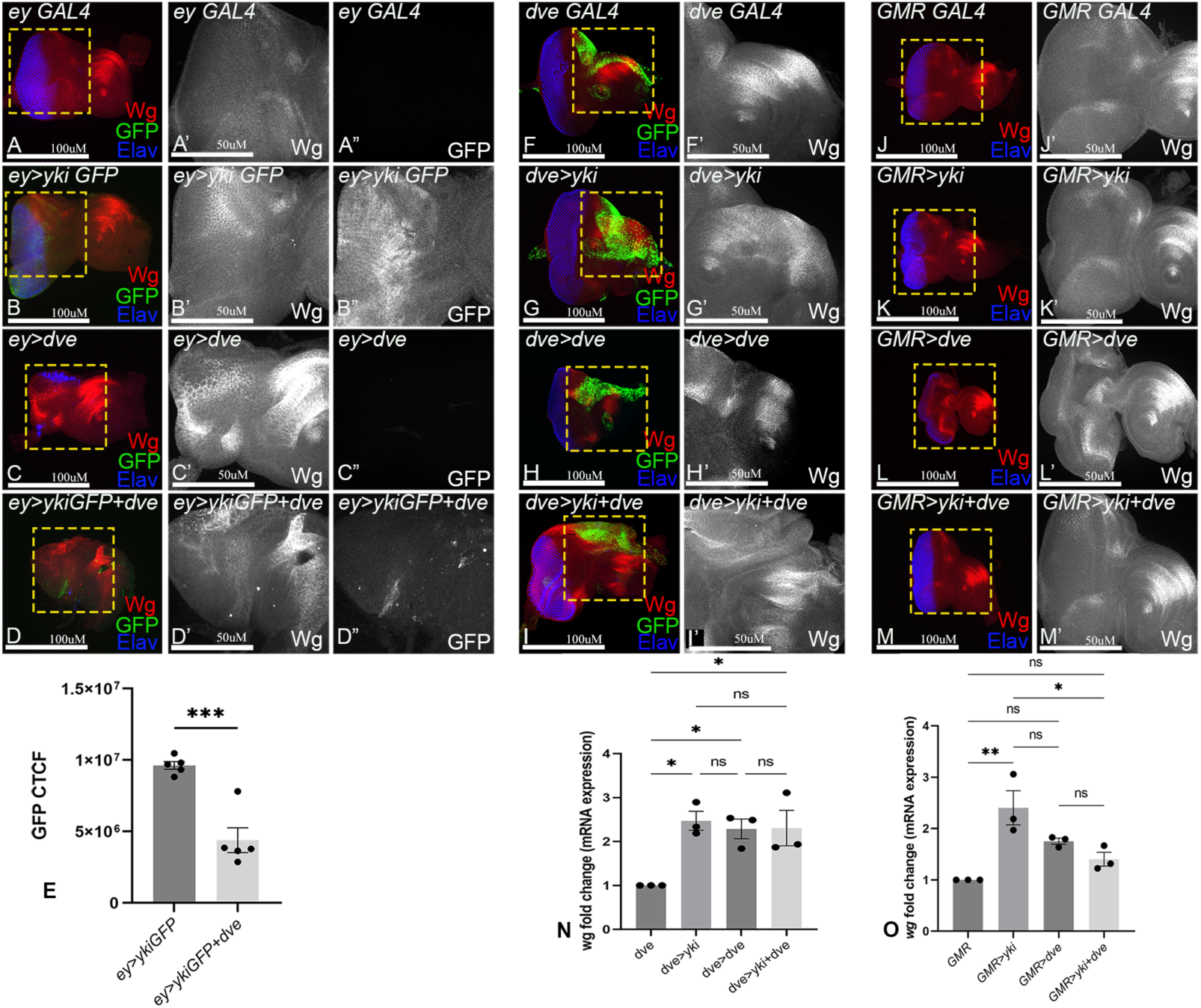
Genetic Epistasis shows Dve acts downstream of Yki. (A-D) Panels show third instar eye discs (A) *ey-GAL4,* (B) *ey-GAL4 UAS yki-GFP* (*ey>yki-GFP*), (C) *ey-GAL4 UAS dve* (*ey>dve*), and (D) *ey-GAL4 UAS yki-GFP, UAS dve* (*ey>Yki-GFP, dve*) showing expression of Wg (red in A-D, grey in A’-D’) and Yki tagged with GFP (green in A-D, grey in A”-D”). Images in A’-D’ and A”-D” are higher magnification images of the box highlighted in A-D. (E) Graph shows quantification of Yki-GFP (CTCF) in the indicated genotypes. (F-I) Panels show expression of Wg (red in F-I, grey in F’-I’) in (F) *dve GAL4 UAS GFP* (*dve>GFP*), (G) *dve GAL4 UAS GFP UAS yki* (*dve>GFP, yki*), (H) *dve GAL4 UAS GFP UAS dve* (*dve>GFP, dve*) and (I) *dve GAL4 UAS GFP UAS yki UAS dve* (*dve>GFP, yki, dve*). Images in F’-I’ are higher magnification images of the box highlighted in F-I. (J-M) Panels show expression of Wg (red in F-I, grey in F’-I’) in (F) *GMR GAL4* (*GMR>*), (G) *GMR GAL4 UAS yki* (*dve>yki*), (H) *GMR GAL4 UAS dve* (*GMR>dve*) and (I) *GMR GAL4 UAS GFP UAS yki UAS dve* (*GMR>yki, dve*). Images in J’-M’ are higher magnification images of the box highlighted in J-M. (N-O) Graphs show quantification of *wg* expression from qRT-PCR when over-expressing Yki or Dve or both with *dve-GAL4* and *GMR-GAL4.* For A,B,C,D,F,G,H,I,J,K,L,M Scale bar = 100µM, and for A’,B’,C’,D’,F’,G’,H’,I’,J’,K’,L’,M’ Scale bar = 50µM.

The *ey-GAL4* driver causes mis-expression of UAS-transgenes very early in the embryonic eye primordium when *ey* expression is turned on (Hazelett et al., 1998). At this early time point transgenes are expressed at the end of embryogenesis in the eye disc primordium prior to axial patterning and MF progression that later delineate patterning and differentiation events. Using *ey-GAL4* (Fig. 5A, S4), we over-expressed a GFP-tagged version of Yki in *ey-GAL4 UAS-yki-GFP* (*ey>yki-GFP* in Fig. 5B), or Dve in *ey-GAL4 UAS-dve* (*ey>dve* in Fig. 5C) or both in *ey-GAL4 UAS yki-GFP, UAS dve* (*ey>yki-GFP, dve* in Fig. 5D, Fig. S4) eye discs, and tested effects on growth and differentiation in mature wandering third instar eye antennal imaginal discs. Compared to control (Fig. 5A), over-expression of Yki-GFP caused over-growth of the eye disc along with suppression of MF in the dorsal and ventral regions resulting in delay in MF progression (Fig. 5B, quantified in Fig. S4A). Over-expression of Dve (Fig. 5C), on the other hand, caused a significant decrease in eye growth and differentiation (quantified in Fig. S4A). Several discs lacked ELAV positive photoreceptors suggesting a complete suppression of MF initiation in these eye discs (Fig. 5C). Interestingly, co-expression of Yki-GFP and Dve (Fig. 5D), showed similar phenotypes in which the growth of the disc was significantly reduced (quantified in Fig. S4A), and no photoreceptor differentiation was observed. Further, Yki-GFP expression was downregulated (Fig. 5D GFP, D” grey). Thus, early in eye development, misexpression of Dve disrupts both the growth and differentiation of eye discs (Fig. S4A) suggesting that Dve may function downstream of Yki.

Next, we tested if a similar effect was observed when the time-window of mis-expression was shifted to second-instar larva stage, when DV patterning occurs. Using *dve-GAL4,* we over-expressed *UAS-yki* (*dve>yki*), *UAS-dve* (*dve>dve*) or both (*dve>yki, dve*) (Fig. 5F-I, quantified in Fig. S4B) and tested effects on overall patterning and growth in mature third instar eye imaginal discs. We observed a high percentage of pupal- lethality in these experiments, for example, overexpression of *yki* showed 28% lethality whereas *dve or yki, dve* overexpression both showed about 38% pupal lethality. Compared to control (Fig. 5F), we observed that over-expression of Yki caused expansion of the *dve* domain (head primordium) and mild suppression of MF progression along the dorsal and ventral margins as reported earlier (Fig. 5G). Over-expression of Dve on the other hand, caused a significant increase in the *dve* domain size and a concomitant decrease in eye growth (Fig. 5H). Interestingly, co-expression of Yki and Dve (Fig. 5I) produced phenotypes reminiscent of Dve-misexpression alone, and the overall Dve domain was reduced compared to Yki overexpression alone (Fig. 5I, quantified in Fig. S4B). Thus, Dve acts downstream of Yki to regulate growth and differentiation.

Next, we tested if a similar effect was observed when the time-window of mis-expression was further shifted to early third-instar larva stage, when the MF is formed, and photoreceptor differentiation begins. Using *GMR-GAL4,* we over-expressed *UAS-yki* (*GMR>yki*), *UAS-dve* (*GMR>dve*) or both (*GMR>yki, dve*) (Fig. 5J-M) and tested effects in mature wandering third instar eye antennal imaginal discs. Compared to control (Fig. 5J), over-expression of Yki caused increased disc size with mild suppression of MF progression in the dorsal and ventral eye disc margins (Fig. 5K), and resulted in 100% viable adult flies at eclosion that show enlarged eyes as previously reported (Wittkorn et al., 2015). Overexpression of Dve caused reduction in imaginal disc growth and suppressed differentiation (Fig. 5L) resulting in formation of smaller eyes with pigmentation defects in adults (quantified in Fig. S4C).

Overexpression of *dve* by *GMR-GAL4* resulted in 87.75% lethality at late pupal stages, while 12.24% of flies that survived at eclosion showed smaller eyes. Interestingly, co-expression of Yki and Dve showed 53.38% pupal lethality and *dve* significantly suppress the enlarged eye phenotype caused by *yki* overexpression (Fig 5M). Taken together, these data support a model in which Dve acts downstream of Yki and negatively regulates Yki activity and its effects on disc growth. Further, using early and later GAL4 drivers, we found that the growth of disc in the early developmental time points (*ey-GAL4*) is weakly suppressed by Dve (Fig. S4A-C). This suggests that Yki is required for early disc growth and proliferation to generate a pool of cells that later undergo differentiation.

In these experiments, we also quantified the expression of Wg, a transcriptional target of both Dve and Yki (Puli et al., 2024; Wittkorn et al., 2015). Interestingly, Yki moderately induced Wg expression when driven by *ey-GAL4* or *dve-GAL4* or *GMR-GAL4* (Fig. 5B,G,K, quantified in Fig. S4D-F) whereas, Wg levels were strongly induced by Dve overexpression in all experiments (Fig. 5C,H,L, quantified in Fig. S4D-F). Consistent with the phenotypes described above, Wg expression was significantly elevated in all samples in which Yki and Dve were co-expressed (Fig. 5D,I,M), further suggesting that Dve may act downstream of Yki in this context. Regulation of Wg was further validated by qRT-PCR (Fig. 5N,O). Compared to control (Fig. 5N,O), overexpression of Yki by *dve-GAL4* (Fig. 5N) or *GMR-GAL4* (Fig. 5O) showed significant induction of *wg* expression. Similarly, overexpression of Dve in its native domain (*dve>dve*) caused significant upregulation of *wg* (Fig. 5N). In contrast, overexpression of Dve outside of its native domain using GMR-GAL4 did not significantly induce *wg* (Fig. 5O) suggesting a spatially restricted effect of Yki-mediated and Dve-mediated *wg* regulation. Together these data indicate that Dve acts downstream of Yki and affects MF progression, dorsal head cuticle versus eye fate by regulating dorsal head cuticle size, and overall patterning of the eye-antennal disc.

### Transcriptional control of DV and Hippo targets during Dve-Yki interactions

The data so far suggest a model where Yki is required for growth and early proliferation of cells in the eye disc, while Dve is expressed in a small group of cells in the dorsal head vertex region and affects both the head cuticle fate and Wg expression. Through these activities, Dve regulates MF progression and the overall number of cells differentiating into photoreceptors. In the eye disc, Yki is ubiquitously expressed and partners with transcription factors such as Sd or Hth/Tsh, whereas Dve is a K50-domain transcription factor that regulates genes in the DV patterning pathway. Given the interactions we observed, we asked if Yki could regulate expression of other genes in the DV patterning pathway, and if Dve could regulate other Yki targets besides Wg. We selected *pannier (pnr)*, an upstream regulator of Dve, and *mirror* (*mirr)* a member of the Iro-C complex that acts downstream of Wg in DV patterning in the eye (Oros et al., 2010; Singh et al., 2012). For assessing Yki targets, we selected *diap1*, an ubiquitously expressed Yki-target gene (Kango-Singh and Singh, 2009). Using qRT-PCR approach, we compared expression levels of these transcriptional targets when Dve or Yki or both were over-expressed in the head cuticle domain defined by *dve-GAL4*, or in the differentiating photoreceptor neurons defined by *GMR-GAL4* (Fig. 6). We found no significant differences in *diap1* expression levels (Fig. 6A, B) when using *dve-GAL4* we overexpressed Yki (*dve>GFP, yki*) or Dve (*dve>GFP, dve*) alone or together (*dve>GFP, yki, dve*) (Fig. 6C). In contrast, using *GMR-GAL4*, *diap1* levels were significantly induced by Yki (*GMR>yki*) but not by Dve (*GMR>dve*). In this domain, Dve did not affect Yki-mediated *diap1* expression (*GMR>yki, dve*) (Fig. 6D). These results indicate that Yki activity is differentially regulated by Dve in a domain dependent manner.

**Figure 6.**
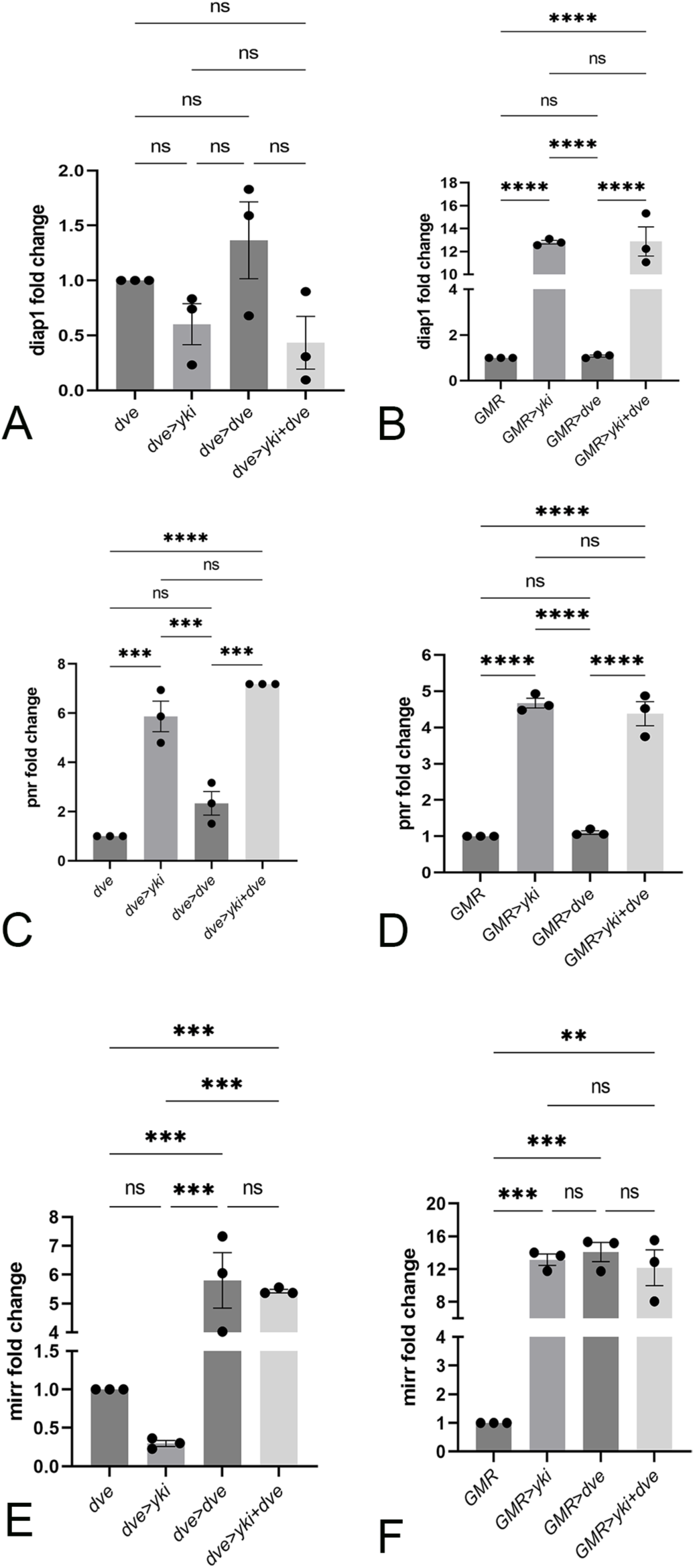
*pannier* and *mirror* are Yki- regulated genes during eye development. RT-qPCR analysis was performed to examine relative changes in expression of (A, B) *diap1*, (C,D) *pannier* and (E,F) *mirror* upon overexpression of Yki or Dve or both, under *dve-GAL4* (A, C, E) or *GMR-GAL4* (B, D, F). Graphs show average fold-changes from three independent experiments, with all samples tested in triplicate for each run. For quantification, bar scatter plots were generated (n=5) and error bars show SEM, P-values are shown as follows: **** p<0.0001, *** p<0.001, * p<0.05, ns., no statistical differences.

We next tested expression of genes in the DV patterning cascade (Singh et al., 2012). We tested the expression of *mirror* (*mirr*) in samples from the *dve-GAL4* set. Compared to control (*dve>GFP*), over-expression of Yki (*dve>GFP, yki*) had no effect on *mirr* expression, whereas over-expression of Dve (*dve>GFP, dve*) caused a significant upregulation of *mirr* expression(Fig. 6E). Expression of *mirr* remained significantly high when Dve and Yki were co-expressed (*dve>GFP, yki, dve*) (Fig. 6E). Thus, in the Dve domain, Dve but not Yki can induce *mirr* expression (Fig. 6E). In contrast using GMR-GAL4, *mirr* expression was induced in all combinations, suggesting that outside of the Dve domain Yki can upregulate *mirr* (Fig. 6F). In DV patterning, *pnr* is expressed in the dorsal eye in the peripodial membrane in a pattern that partially overlaps Dve, and acts upstream of Dve (Puli et al., 2024). Interestingly, Yki overexpression robustly induced *pnr* expression using either *dve-GAL4* (*dve>GFP, yki*, Fig. 6C) or *GMR-GAL4* (*GMR>Yki*, Fig. 6D). Overexpression of Dve did not affect *pnr* expression in both experimental settings (Fig. 6C, D), and co-expression of Dve with Yki did not modify Yki-mediated upregulation of *pnr* (Fig. 6C, D). This result suggests that *pnr* is a new Yki target gene. Taken together, these data indicate the Yki can transcriptionally induce multiple genes like *pnr, wg* and *mirr* in the DV patterning hierarchy including *pnr, wg* and *mirr*.

To test if *pnr* in turn affects Yki-Dve interactions, we tested dominant genetic interactions between Yki and *pnr*. Compared to control (*dve>GFP*) (Fig. 7A, quantified in Fig. 7E), loss of one copy of *pnr* (*dve>GFP; pnr^VX6^/+*) did not modify the size of the *Dve-GAL4* expression domain (Fig. 7B, quantified in Fig. 7E). However, the increase in *dve* domain size by Yki overexpression (*dve>GFP, yki*, Fig. 7C, quantified in Fig. 7E) is significantly reduced by loss of one copy of *pnr* (*dve>GFP,yki; pnr^VX6^/+*) (Fig. 7D, quantified in Fig. 7E). These results suggest that Yki and Pnr genetically interact, and Yki activity is differentially regulated the dorsal head cuticle domain versus in the photoreceptor cells. Further, these findings indicate that maintaining appropriate Pnr levels is important for dorsal identity, DV patterning, and proper growth of the eye disc (Fristrom and Fristrom,; Kumar, 2011). Taken together, rescue of *pnr* loss of function by Yki gain of function supports an epistatic relation where *pnr* acts downstream of Yki and regulates Dve expression/function.

**Figure 7.**
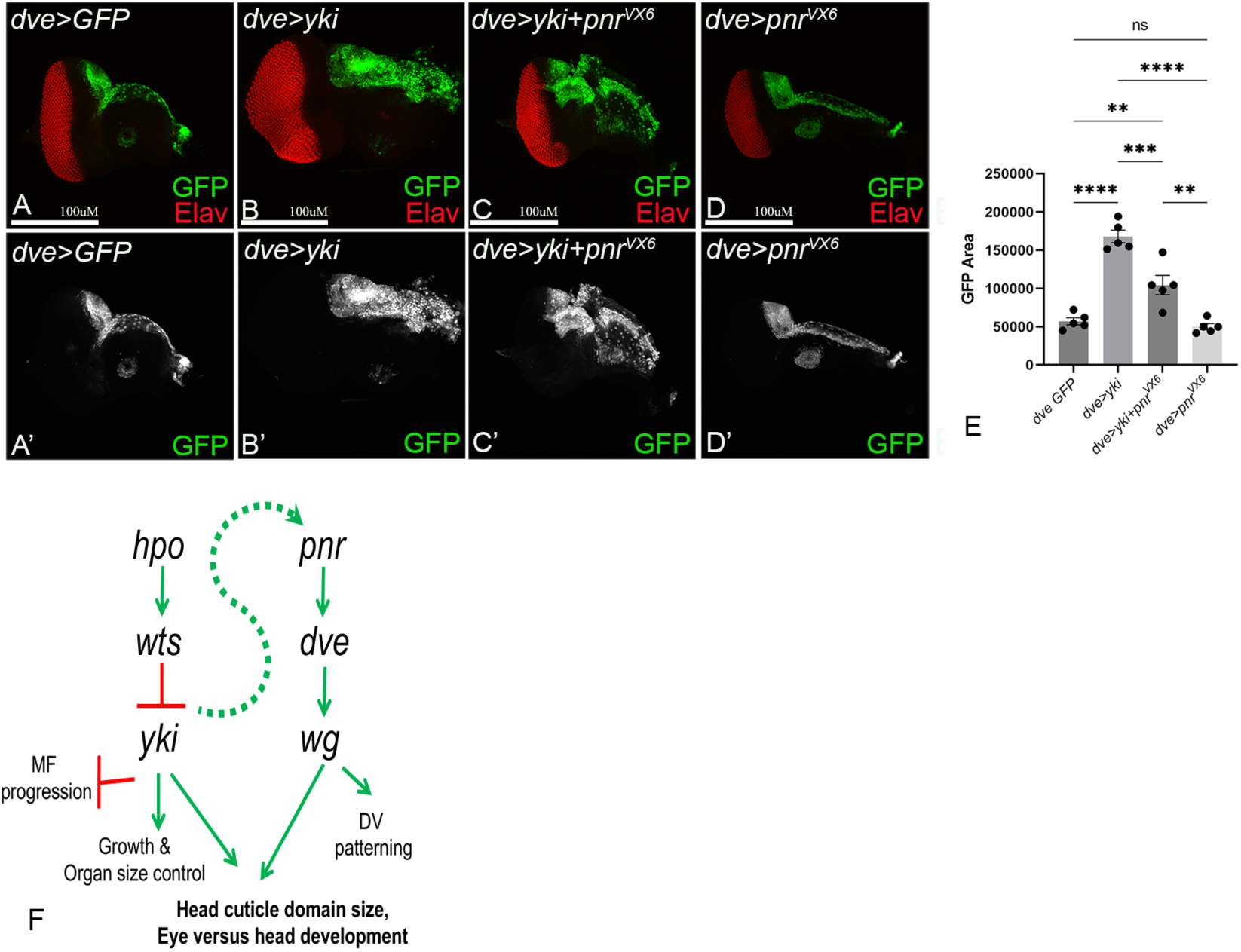
Yki genetically interacts with *pnr*, a DV patterning genes. Panels show confocal images of third instar eye imaginal discs stained for ELAV (red in A-D, grey in A’-D’) from larvae of the following genotype: (A) *dve-GAL4 UAS GFP* (*dve>GFP*) control; (B) *dve-GAL4 UAS GFP UAS yki* (*dve>GFP, yki*), (C) *dve-GAL4 UAS GFP UAS yki FRT82B pnr^VX6^/+* (*dve>GFP, yki, pnr^VX6^/+*) and (D) *dve-GAL4 UAS GFP FRT82B pnr^VX6^/+* (*dve>GFP, FRT82B pnr^VX6^/+*). (E) Quantification of head domain size based in GFP expression is shown in the graph. Scale bar = 100µM. (F) Model proposed for the interactions of Hippo and DV patterning pathways. Independent and shared functions of the two pathways are shown. During development and patterning of the eye imaginal disc, the Hippo pathway effector Yki interacts with *pnr* and *dve* in the DV patterning pathway to promote head cuticle domain size.

## Discussion

Regulation of organ size is controlled by coordination of cell proliferation, survival and growth. The *Drosophila* eye imaginal disc is an elegant model for studying the coordination of growth with differentiation during development due to the well-documented regulation of cell proliferation, developmental apoptosis and patterning from embryogenesis to adulthood (Borday et al., 2012; Kumar, 2023; Singh et al., 2012; Tare et al., 2016; Weasner and Kumar, 2022a). Eye discs are specified in late embryogenesis and expand during larval stages, reaching ∼50,000 cells by the mature third instar, when photoreceptors differentiate posterior to the MF and anterior cells give rise to the head cuticle, antenna, and associated structures (Bessa et al., 2002; Haynie and Bryant, 1986; Ma and Moses, 1995; Tare et al., 2016). During pupal development, extensive tissue remodeling—including head eversion and formation of adult head structures—further refines these domains (Haynie and Bryant, 1986). Notably, both the mature third instar and early-to-mid pupal stages are especially sensitive to developmental perturbations that can manifest as overt morphological abnormalities in the adult.

During organogenesis, successful patterning of the eyes, ocelli, antennae, and maxillary palps requires precise integration of multiple signaling pathways, including Wg, Decapentaplegic (Dpp), Hedgehog (Hh), the retinal determination network, and growth-regulatory mechanisms such as the Hippo pathway. Specifically, Yki functions in early retinal differentiation (Wittkorn et al., 2015), and as shown in this study, in regulating dorsal head domain size via Dve interactions. Our findings thus illustrate how a single transcriptional co-activator can drive distinct outcomes by engaging with different molecular partners. In the context of eye development, DV patterning is the first lineage restriction event controlled by the expression of Pannier (Pnr) which lies upstream of the DV patterning hierarchy. We recently showed that Pnr acts upstream of and regulates the expression of the K-50 domain transcription factor Dve (Puli et al., 2024). Thus, *dve* and *pnr* can have some overlapping functions in dorsal fate determination but Dve mediated Wg regulation is different from Pnr (Puli et al., 2024). Further, Pnr driven Wg regulation in dorsal eye and head region does not represent the complete spectrum of its function (Puli et al., 2024). Specifically, the current study reveals that Yki interacts with Pnr to regulate dorsal head cuticle development, whereas it influences the morphogenetic furrow progression and retinal differentiation by interacting with Wg. Loss of *dve* or other dorsal genes (e.g., *pnr* or Iro-C complex) in the DV patterning pathway cause dorsal eye enlargements due to increased growth (Fristrom and Fristrom,; Kumar, 2011; Maurel-Zaffran and Treisman, 2000; Oros et al., 2010; Singh et al., 2012). Gain of function of Dve or Pnr results in reduced growth and smaller head/eye domains. Similar effects on organ size are well-documented for the Hippo pathway, where loss of function of Hippo signaling results in enlargement of structure due to over-growth caused by increased proliferation and reduced apoptosis; while gain-of-function of Hippo signaling results in formation of smaller structures due to increased apoptosis (Kango-Singh and Singh, 2009). Here, we report that in the context of the eye imaginal discs, the Hippo pathway regulates the head cuticle domain (Fig. 1-2, Fig. S1) in addition to its previously reported role in regulating morphogenetic furrow progression (Peng et al., 2009; Wittkorn et al., 2015). Further, we show that loss of function of Hippo pathway results in expansion of Dve expression and the head cuticle domain associated with the distinct ‘large-headed’ phenotype (Fig. S1C-E) identified through mosaic genetic screens that uncovered upstream components of the Hippo pathway (Kango-Singh and Singh, 2009; Verghese et al., 2020). Thus, both the DV patterning and Hippo pathway genes play a significant role in organogenesis of the head.

In *Drosophila*, eye fate is antagonistic to the fates of other domains in the eye antennal disc during patterning, as seen by antagonistic interactions between eye and antenna fate (Lengler, 2001), and Pnr-regulated dorsal eye margin and eye fate (Singh et al., 2006; Singh et al., 2011). Similar antagonistic interactions are evolutionarily conserved in mammalian models. For example, antagonistic interactions between Wnt and Hh pathway, and TGFb and BMP pathways regulate retinal development and differentiation. Similarly Srbep2 and Lrp2 regulate eye size, and Six3 and Prox1 regulate lens development by acting antagonistically (Koontz et al., 2013; Mai et al., 2022; Ulloa and Martí, 2010). Together, these examples illustrate that antagonistic interactions between developmental signaling pathways are evolutionarily conserved mechanisms for head and eye development across species.

In addition to the developmental patterning pathways, imbalance in the activity of growth regulatory pathways also differentially affects eye versus head cuticle fate. Activation of Yki either through over-expression of *yki^3SA^*, or loss of *hpo* or *wts*, leads to expansion of the head cuticle domain and suppression of eye fate (Fig. 2, Fig. S1).

Interestingly, this effect is suppressed by overexpression of Dve, suggesting an antagonistic interaction between Yki and Dve (Fig. 1). Over-expression of Dve suppresses Yki activity as seen by downregulation of DIAP1 (Fig. S2). Further, Sd also shows antagonism to Dve regulated head cuticle domain (Fig. S1). Thus, genetic interactions studies place Dve downstream of Yki, functioning as a suppressor of Yki activity.

Furthermore, the role of Dve in delimiting spatial domains (eye versus antenna or eye versus head) produces interesting phenotypes that extend well beyond the Dve expressing dorsal head vertex domain. For example, we found that Dve exerts differential effects on Yki activity, depending on the spatial context. Over-expression of Dve within its endogenous domain resulted in suppression of Yki activity as seen by altered expression of Hth (Fig. 4), whereas expression of other Yki targets, like *ex* or DIAP1, was not significantly affected (Fig. 3, Fig. S2). These data suggest that boundary-dependent or antagonistic interactions determine a balance between activation and suppression of Hippo pathway coactivator in *dve*-expressing versus non-expressing cells. This spatially restricted regulation results in differential Dve-mediated regulation of Yki activity. Together, these interactions play a major role in regulating optimal eye and head size during development.

In a model of wound induced regeneration in wing discs, *dve* is upregulated by *Y*ki overexpression and downregulated by *yki* knockdown, suggesting that *dve* is a target of Yki (Zhang et al., 2017). Consistently, RNAseq data identified *dve* as a potential Yki target in wing discs (Zhang et al., 2017). Yki associates with some of its transcriptional targets through WW-PPxY interactions, for example, the PPxY domain of Ex associates with the WW-domain of Yki (Guo et al., 2025; Kango-Singh and Singh, 2009). However, Dve lacks a PPxY motif suggesting that it does not directly interact with Yki via this mechanism. As a transcriptional co-activator Yki lacks a DNA-binding domain and instead associates with sequence specific DNA-binding transcription factors to regulate downstream targets. In the eye-antennal disc, Yki partners with transcription factors like Sd which is expressed in the photoreceptor domain (Goulev et al., 2008; Wu et al., 2008), and Hth or Tsh, which are expressed anterior to the morphogenetic furrow and in the antenna, ptilinum, and ocellus. These factors define the boundary of the eye field and the head capsule in the eye disc (Bessa et al., 2002; Lopes and Casares, 2010; Singh and Choi, 2003; Wang and Sun, 2012). Using a *dve-lacZ* reporter (Nakagoshi et al., 1998) and qRT-PCR analysis, we found that *dve* is not directly regulated by Yki at the transcription level during eye development (Fig. S3). These data suggest that *dve* regulation of Yki is tissue- and context-specific, while *dve* is transcriptionally regulated by Yki during wing disc regeneration, dve is not regulated by Yki during patterning and growth of eye antennal disc during development.

The development of head cuticle is a critical process for survival after metamorphosis. DV patterning requires sequential and regulated expression of several genes in cells that differentiate into the head cuticle, ocelli and antennae. Several signaling pathways like Hedgehog, Notch, and DV (Pnr/Dve) patterning during larval stages (Chang et al., 2001; Kango-Singh and Singh, 2009; Singh et al., 2012), and Osiris gene family during pupal stages (Sun et al., 2024a), play essential roles in ensuring normal head development. These pathways act in coordination with growth-regulatory pathways throughout development. Yki partners with Sd or Hth or Tsh to regulate growth-promoting activities by controlling the expression of target genes involved in organ growth, cell proliferation (e.g., *cyclin E*, *cyclin B*, others), and cell survival (*diap1*, *bantam miRNA*). Hth maintains proliferation of uncommitted retinal progenitor cells in the eye disc (Lopes and Casares, 2010), and together with transcriptional partners like Yki regulates *miRNA bantam* and *dpp* expression (Ikmi et al., 2014; Peng et al., 2009; Slattery et al., 2013). Yki can also partner with other transcription factors to regulate diverse biological processes, for example, Foxo in response to oxidative stress (Sun et al., 2024b), and GAGA factor (GAF) and Brahma (BRM) complex for chromatin remodeling (Oh et al., 2013). Further, transcriptional repressor proteins like E2F (Zhang et al., 2017) or Tondu-domain-containing growth inhibitor (Tgi) (Guo et al., 2025) can compete with Yki for binding with Sd thereby modulating Yki transcriptional activity. Mammalian YAP and TAZ also bind to several transcription factors (e.g., TEAD1-4, p73, Runx2, β-catenin, SMADs) to regulate growth-promoting and oncogenic activities(Hong and Guan, 2012; Zhao et al., 2008). The large repertoire of genes regulated by Yki/YAP in multiple biological contexts led us to ask if additional genes within the DV patterning hierarchy might be regulated by Yki. Using genetic interactions (Fig. 5, Fig. 7) and qRT-PCR assays (Fig. 6), we found that *pnr* and *mirr* (member of the Iro-C complex) are transcriptionally regulated by Yki. In addition, *wg* is a shared target of both Yki and Dve during the growth and patterning of the eye discs (Fig. 5, Fig. S4). Taken together, our data show multiple regulatory interactions between the Hippo pathway effector Yki and components of the DV patterning pathway that are essential for coordinated growth and patterning of the eye and head in flies.

The interactions among *yki, dve, pnr* and *wg* are particularly significant given the conserved roles of their mammalian orthologs YAP1/2, SATB1/2, GATA3/4/5 and Wnt5 in development and disease (Bai et al., 2025; Bejsovec, 2018; Hellebrekers et al., 2009; Nishioka et al., 2009; Ralston et al., 2010; Waghmare et al., 2023). YAP–TEAD–GATA3 signaling is essential for trophoblast and trophectoderm specification across species, and aberrant YAP activity is associated with defects in early embryonic development (Gerri et al., 2020; Lin et al., 2023). SATB1 and SATB2 regulate multiple developmental processes, including neurogenesis and craniofacial patterning, and mutations in these genes result in severe neurodevelopmental and craniofacial abnormalities like dysmorphic facial features including brachycephaly (a flat head shape), cleft palate, prominent forehead, hypertelorism (widely spaced eyes), and low nasal bridge (Agrelo et al., 2009; Kuo et al., 2025; Robinson et al., 2025; Savarese et al., 2009; Wahl et al., 2024, Britanova et al., 2005; Dobreva et al., 2006; Naik and Galande, 2019). Both factors are also implicated in tumor progression, where elevated SATB1/2 expression correlates with aggressive behavior and metastasis (Roy et al., 2020). Moreover, YAP–SATB2 interactions have been documented in cortical neuron development and in renal cell carcinoma, where YAP/TEAD4 drives SATB2-dependent proliferation (Naik and Galande, 2019; Roy et al., 2020, Su et al., 2022, Jin et al., 2023). Together, these conserved relationships highlight the relevance of our findings and provide a framework for exploring how growth and patterning pathways integrate to regulate head development. Further elucidation of the molecular mechanisms underlying these interactions will refine our understanding of coordinated tissue growth across species.

## Materials and Methods

### Drosophila stocks

All fly stocks used in this study are described in Flybase (www.flybase.org): *w, bi-GAL4* (Lecuit et al., 1996), *w; dve-GAL4* (Nakagoshi et al., 1998), *w; ey-GAL4* (Hazelett et al., 1998), *w; GMR-GAL4* (Moses and Rubin, 1991), *yw; FRT42D dve^1/^CyO* (Nakagoshi et al., 1998), *yw; FRT42D hpo^42-47^/CyO* (Wu et al., 2003), *eyFLP; FRT42D cl w^+^/ CyO, y^+^* (BDSC # 5617), *w; +; FRT82B wts^X1^/TM6B* (BDSC# 44251), *yw eyFLP; FRT82B cl w^+^*/*TM6B* (BDSC# 43657), *yw; FRT82B pnr^VX6^/TM6B* (BDSC #6334), *w; UAS GFP* (BDSC# 4775), *w; UAS dve* (Nakagoshi et al., 1998), *w; UAS dve^RNAi^* (Nakagoshi et al., 1998), *w; UAS hpo* (Udan et al., 2003), *w; UAS wts^13F^* (Kwon et al., 2015), *w; UAS yki* (Huang et al., 2005), *w; UAS yki-GFP* (Oh and Irvine, 2009)*, w; UAS yki^3SA^* (Oh and Irvine, 2009), *w; UAS yki^RNAi^* (BL31965), *w; UAS sd* (BDSC# 9373); *w; UAS sd^RNAi^* (BDSC #32934), *w; dve^1^-lacZ/Cy*O (BDSC# 11073), *w; ex^697^-lacZ/CyO* (Boedigheimer and Laughon, 1993), and *w; bantam-sensor GFP* (Thompson and Cohen, 2006).

### Immunohistochemistry

Third instar eye imaginal discs were dissected in PBS and fixed in 4% paraformaldehyde fixative for 20 min. Fixed discs were washed in 0.3%PBTX for 10min, three times and blocked in 2% PBTN (0.3% PBTX and 2% Normal Donkey Serum). The following primary antibodies were used: Rat anti-Elav (7E8A10, 1:100), mouse anti-Wg (24D10, 1:100), mouse anti-βGAL (40-1a, 1:100) all from DSHB, Iowa, IA, USA, and mouse anti-DIAP1 (1:200, a gift from Dr. Bruce Hay). Rabbit anti-Dve (1:1000), and rabbit anti-Hth (1:250) are polyclonal affinity purified antibodies reported in this study (ThermoFisher). The secondary antibodies were donkey Fab fragments conjugated to FITC, Cy3 or Cy5 against rabbit, mouse, or rat hosts (Jackson ImmunoResearch Labs, West Grove, PA, USA).

### Image acquisition and analysis

#### Confocal microscopy

The eye antennal discs were imaged with a confocal laser scanning microscope (Olympus FV3000) with FV10-ASW 4.0 software, at 1024x1024 pixels per image, using Plan-Apochromat 20X and 40X objectives.

### Bright field microscopy

Adult eyes were imaged at 10X using MrC5 color camera mounted on a Axioimager.Z1 Zeiss Apotome using the *z*-sectioning function of Axiovision software 4.6.3. Adult flies were frozen at −20°C for ∼4h and mounted on a needle for imaging. All imaginal discs, and adult flies are imaged at identical magnification, and IHC stains imaged under identical confocal settings. The images were further processed and analyzed using Adobe Photoshop 2025 and Fiji/ImageJ (NIH). The disc orientation is dorsal up and anterior to the right in all figures.

### Image analysis

The Z stacks obtained from confocal imaging were analyzed by Fiji/ImageJ software to quantify area, and fluorescence intensity. Area of selected regions of interest (ROI) were outlined by referring to GFP-positive regions to mark Dve expressing cells and Elav positive cells to mark differentiating photoreceptor cells across the entire eye disc. Area and fluorescence measurements were obtained by using the ‘Measure’ tool. Total fluorescence was calculated using the formula in the Image J program, CTCF (Corrected Total Cell Fluorescence) **=** (Integrated Density of ROI) **–** (Area of ROI X Mean fluorescence of background readings) (https://theolb.readthedocs.io/en/latest/imaging/measuring-cell-fluorescence-using-imagej.html;(Rohith and Shyamala, 2017)). Integrated density was calculated as Mean Intensity X Area of the selected ROI.

### Statistical Analyses

Statistical analyses were performed using GraphPad Prism (v9.5) software. One-way ANOVA was applied, followed by post-hoc analyses using Tukey’s test for comparisons among multiple groups, and a *t*-test was used for comparisons between two groups. Graphs are presented as bar scatter plots showing mean ± SEM. Five eye-antennal imaginal discs (n=5) were subjected to quantification and analyzed for each control and experimental group. Asterisks indicate significant differences based on *p*-values: ns = *p* > 0.05, *= *p* < 0.05, **= *p* < 0.01, ***= *p* < 0.001.

### qRT-PCR and quantification

Eye imaginal discs were dissected from control and experimental groups, stored in TRIzol at -80°C, and total RNA was extracted using the ZYMO RNA Clean and Concentrator Kit (Cat No.# R1050). A total of 500ng RNA was used for cDNA synthesis using the First Strand cDNA synthesis kit (Cytiva, Danaher Corporation, Washington, DC, USA, Cat No.#27926101) following the manufacturer’s guidelines. qRT-PCR was performed using iQ-SYBER Green Supermix (Bio-Rad laboratories, Hercules, CA, USA; Cat No.# 1708882) with each sample in triplicate on the iCycler iQ™ Real-Time PCR Detection System (Bio-Rad Laboratories, Hercules, CA, USA). Each primer set was tested in three independent runs (three technical replicates), and relative gene expression was calculated using the ΔΔCT method (2−ΔΔCT method). Statistical analyses of the qRT-PCR data were performed using GraphPad Prism (v9.5) using the Ordinary One-way ANOVA method, followed by correction for multiple comparisons using the Tukey’s post-hoc test assuming statistical significance at *p≤0.05*. Graphs were generated using GraphPad Prism (v9.5).

The following primers were used:

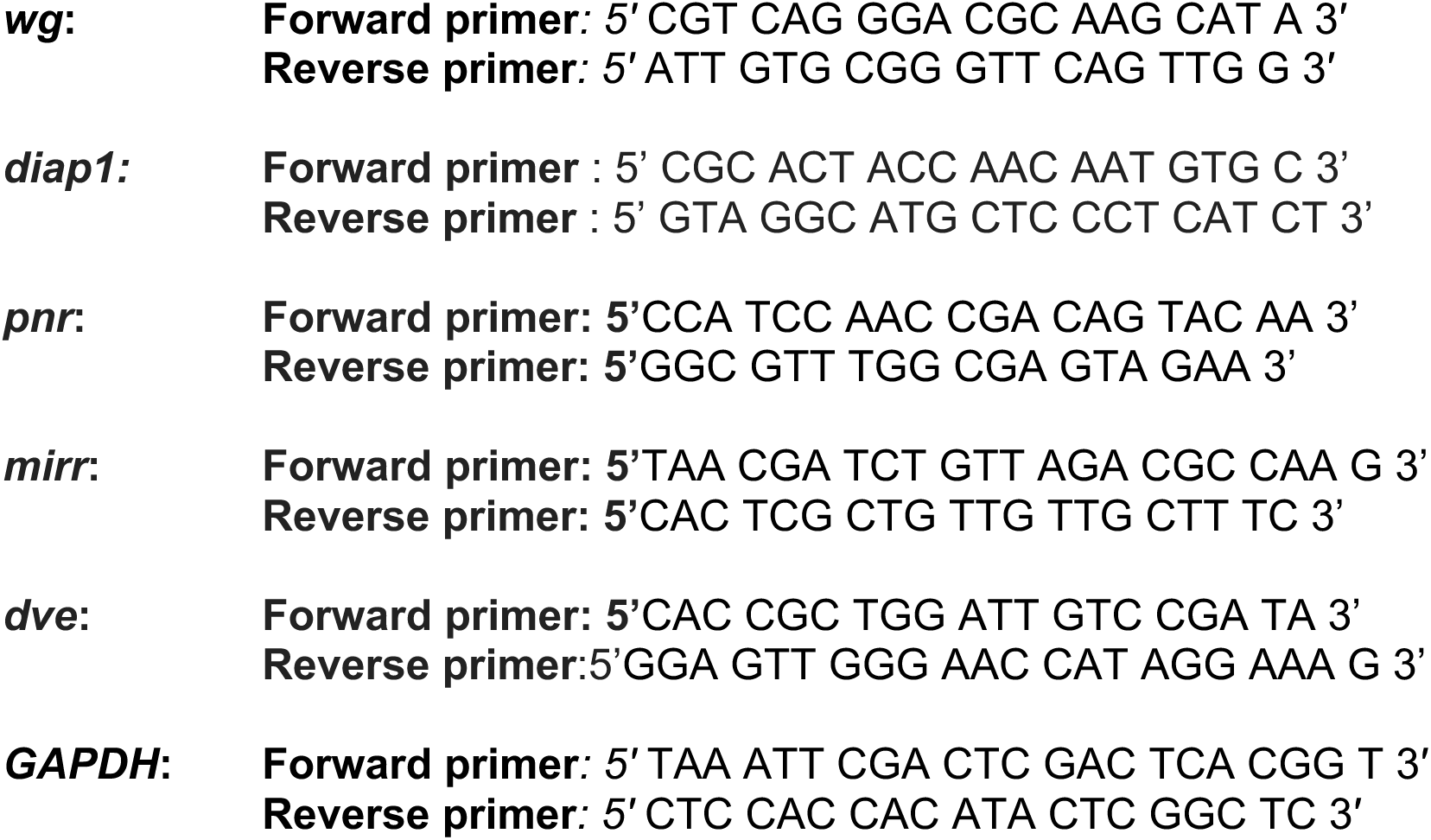

## Supporting information

Figure S1 (linked to Figure 2): Modulating Hippo activity affects head cuticle domain size.

Supplementary Figure S2 (linked to Figure 3): Dve modulation and its effects on Yki activity.

Supplementary Figure S3 (linked to Figure 4): Yki does not transcriptionally regulate dve.

Supplementary Figure S4 (linked to Figure 5): Effects on eye disc area and Wg expression in genetic epistasis interactions.

## Acknowledgments

The authors would like to thank the members of the Kango-Singh and Singh laboratory for their support and helpful discussions. For monoclonal antibodies used in this study, we thank the Developmental Studied Hybridoma Bank (DSHB), created by the NICHD of the NIH and maintained at The University of Iowa, Department of Biology, Iowa City, IA 52242. Stocks obtained from the Bloomington Drosophila Stock Center (NIH P40OD018537) were used in this study. Work in MKS lab and AS labs is supported by the National Institutes of Health (1R01EY032959-01, MPIs AS and MKS). MKS lab is also supported by funds from the Schuellein Chair in Biological Sciences, and work in AS lab is supported by Leonard Mann Chair in Natural Sciences. AR and NG were supported by the Graduate Teaching Assistantship from University of Dayton.

## Author Contributions

BNR and NG performed experiments, BNR, AR, NG did image acquisition and analyses, and quantification of data, MKS and AS obtained funding, MKS oversaw the project and wrote the manuscript with inputs from all authors.

---

**Figure S1 (linked to Figure 2):**
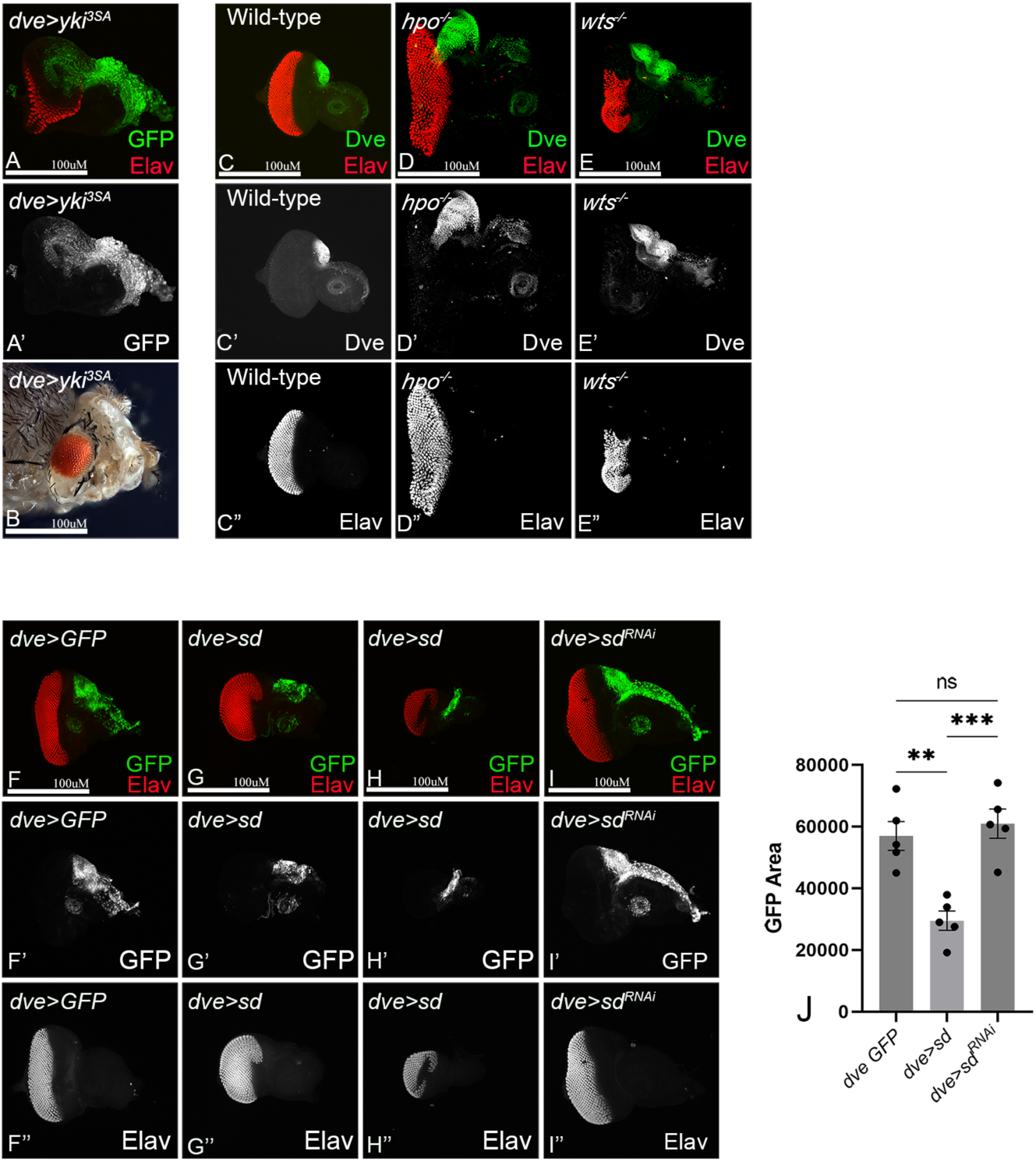
Modulating Hippo activity affects head cuticle domain size. (A-B) Panels show third instar eye imaginal disc (A) and the corresponding adult (B) from *dveGAL4>UAS GFP, UAS yki^3SA^* showing effects of hyperactivation of Yki on eye (A), head cuticle size (A’), and adult (B). (C-E) Panels show third instar eye imaginal discs from (C) wild type, (D) *FRT42D hpo^42-47^/FRT 42D cl w^+^,* and (E) *FRT82B wts^X1^/FRT82B cl w^+^* showing expression of ELAV (red in C-E, grey in C”-E”) and Dve (green in C-E, grey in C’-E’). (F-J) Panels show third instar eye discs (F) wild-type, (G-H) *dve-GAL4> UAS GFP, UAS sd* (*dve>GFP, sd*), and (I) *dve-GAL4> UAS GFP, UAS sd^RNAi^*(*dve>GFP, sd^RNAi^*) showing expression of ELAV (red in F-I, grey in F”-I”) to measure eye size. The *dve* expression domain size was monitored by GFP expression (green in F-I, grey in F’-I’). (J) Quantification of the head domain size based on GFP expression is shown in the graph. Scale bar = 100µM.

**Supplementary Figure S2 (linked to Figure 3):**
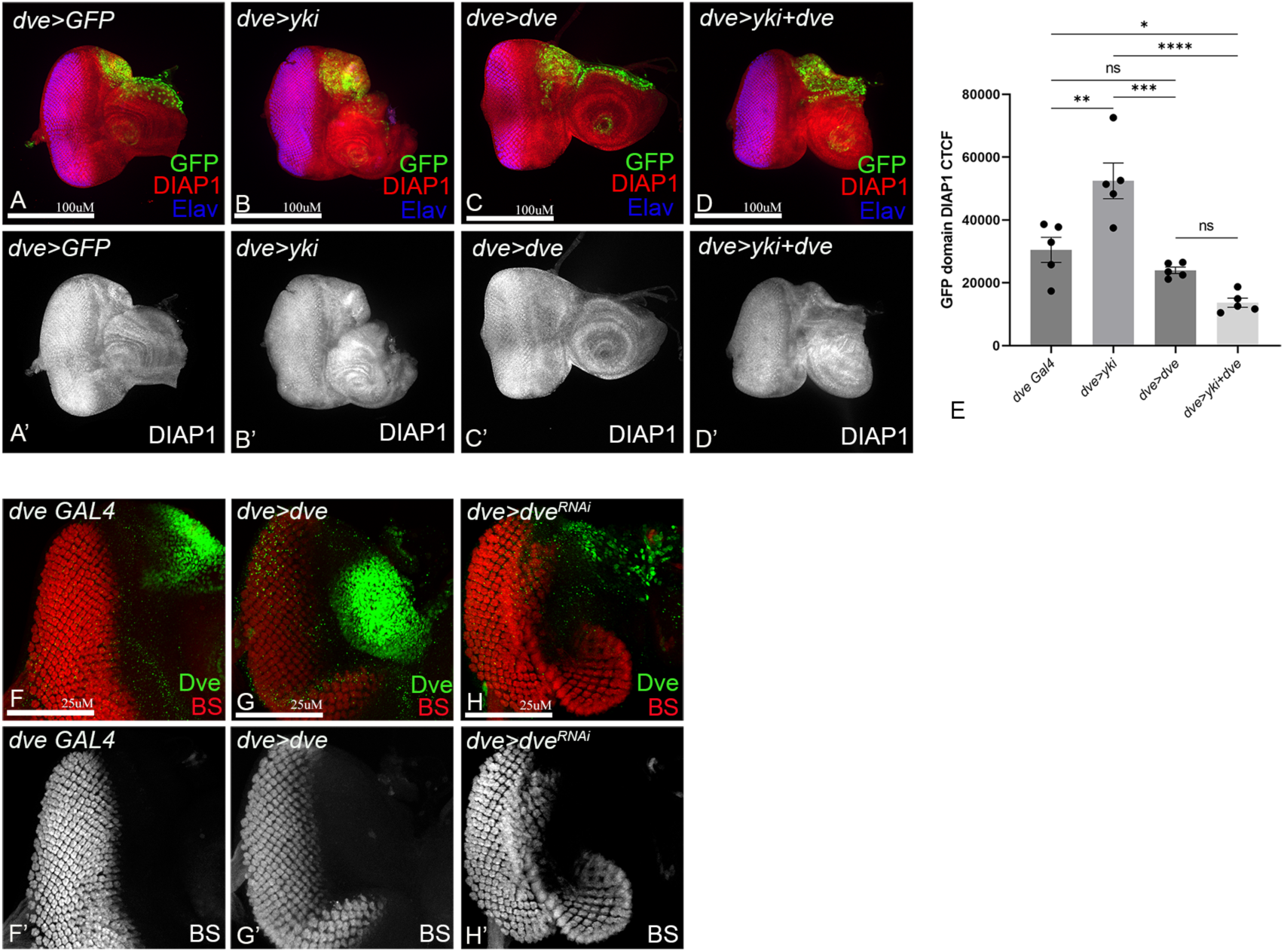
Dve modulation and its effects on Yki activity. (A-E) Confocal images of third instar eye imaginal discs stained for DIAP1 (red in A-D; grey in A’-D’) from (A) *dve-GAL4>UAS GFP* control *(dve>GFP)*, (B) *dve-GAL4 UAS GFP UAS yki* (*dve>GFP, yki*), (C) *dve-GAL4 UAS GFP UAS dve* (*dve>GFP, dve*) and (D) *dve-GAL4>UAS GFP UAS yki UAS dve* (*dve>GFP, yki, dve*). (E) Quantification of Diap1 expression (CTCF) is shown in the graph. (F-H) Panels show confocal images of eye discs showing expression of the *bantam sensor-GFP* (red in F-H, grey in F’-H’) and Dve (green in F-H) from (F) *dve-GAL4 bantam-sensor-GFP* (*BS-GFP, dve>*) control, (G) *dve-GAL4 bantam-sensor-GFP UAS dve* (*BS-GFP, dve>dve*), and (H) *dve-GAL4 bantam-sensor-GFP UAS dve^RNAi^* (*BS-GFP, dve>dve^RNAi^*). For A,B,C,D, Scale bar = 100µM. For F,G,H Scale bar = 25µM.

**Supplementary Figure S3 (linked to Figure 4):**
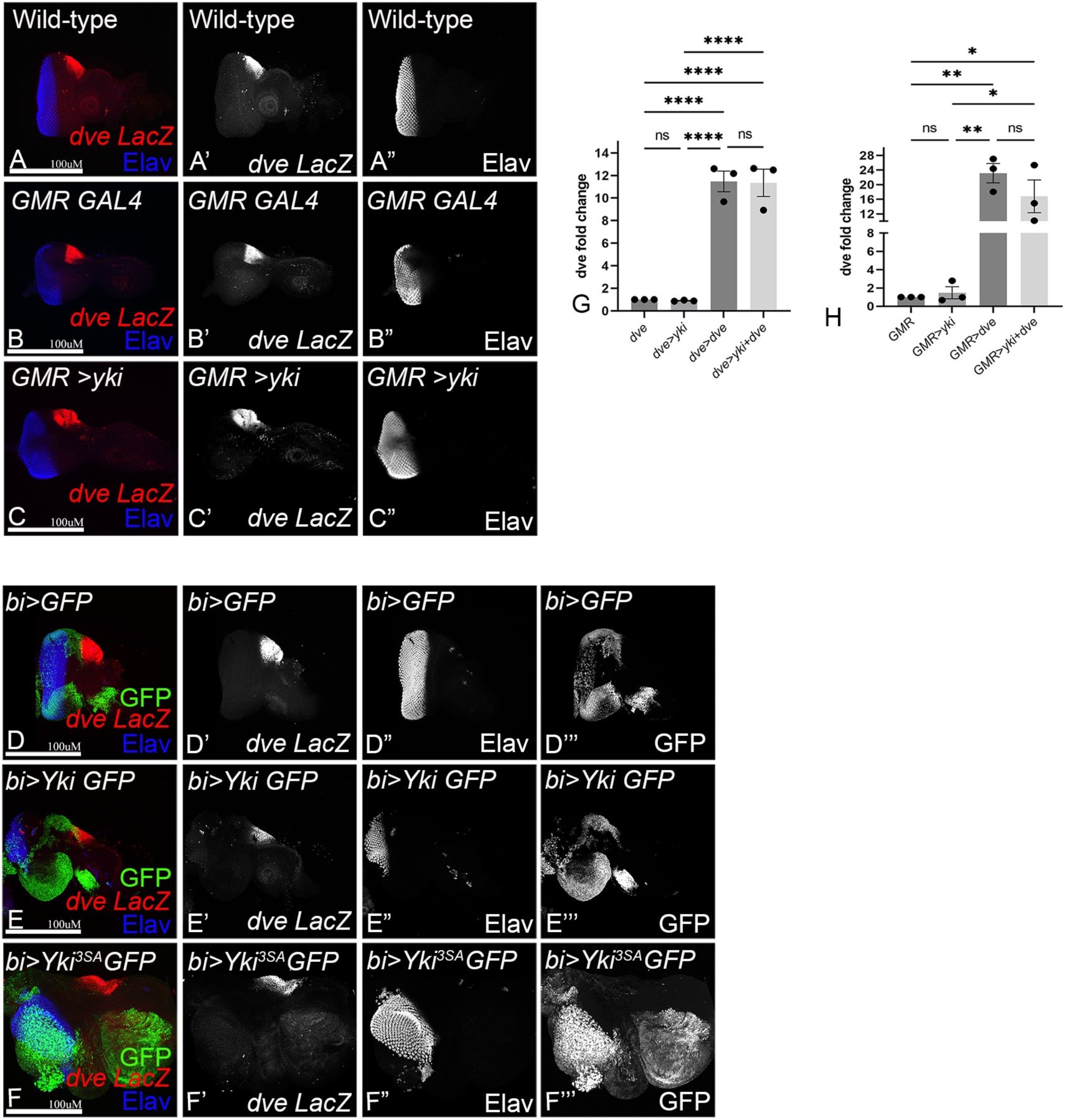
Yki does not transcriptionally regulate *dve*. (A-C) Panels show confocal scans of third instar eye discs showing expression of the *dve-lacZ* reporter (red in A-C, grey in A’-C’) and ELAV (blue in A-C, grey in A”-C”) from the following genotypes: (A) Wild-type (B) *GMR-GAL4* (*GMR>)* control, (B) *GMR-GAL4 UAS yki* (*GMR>yki*). (D-F) Panels show confocal scans of third instar eye discs showing expression of *dve-lacZ* (red in A-C, grey in A’-C’) and ELAV (blue in A-C, grey in A”-C”) from (D) *bi-GAL4 UAS GFP* (*bi>GFP*) (E) *bi-GAL4* UAS GFP, UAS yki (*bi>GFP, yki*) control, (F) *bi-GAL4 UAS GFP UAS yki^3SA^* (*bi>GFP,yki^3SA^*). (G. H) Graphs showing quantification of *dve* expression from qRT-PCR upon over-expression of Yki or Dve or both using *dve-GAL4* and *GMR-GAL4.* Graphs show average fold-change from three independent experiments wherein all samples were tested in triplicate for each run. Scale bar = 100µM.

**Supplementary Figure S4 (linked to Figure 5):**
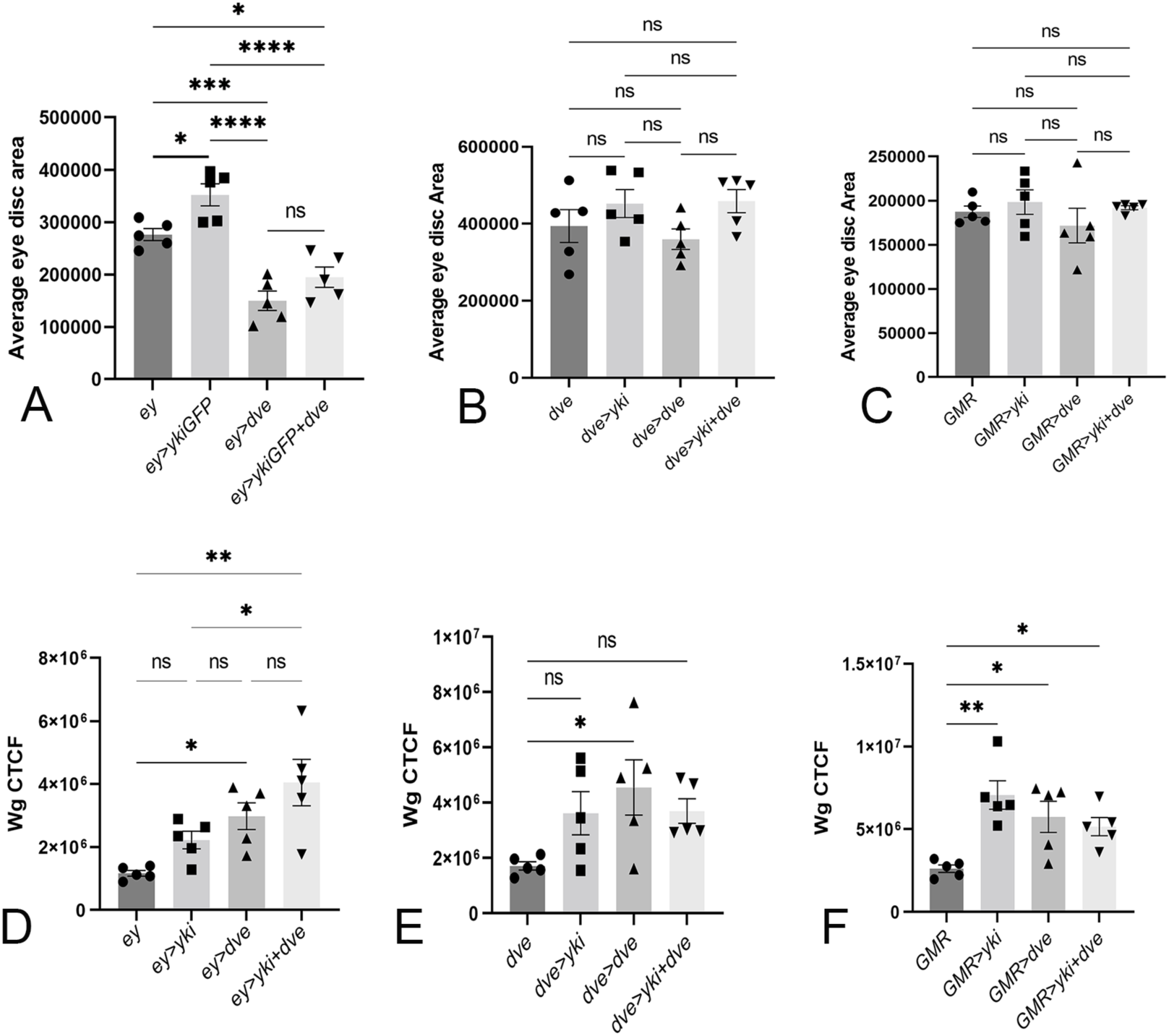
Effects on eye disc area and Wg expression in genetic epistasis interactions. Quantification of average eye disc area following overexpression of either Yki or Dve or both using (A) *ey-GAL4*, (B) *dve-GAL4 UAS GFP*, and (C) *GMR-GAL4*. (D-F) Graphs showing quantification of *wg* expression (CTCF) upon over-expression of Yki or Dve or both with *ey-GAL4*, *dve-GAL4* and *GMR-GAL4.* Graphs in D-F show average fold-changes from three independent experiments, with all samples tested in triplicate for each run. For quantification, bar scatter plots were generated (n=5), and error bars represent SEM, P-values are shown as follows: **** p<0.0001, *** p<0.001, * p<0.05, ns., no statistically significant differences.

