## Supplementary figures and images for "A Genetic Mechanism Linking Hippo Signaling to Dorsoventral Patterning for Control of Head and Eye Development"

### Figure S1 (linked to Figure 2): Modulating Hippo activity affects head cuticle domain size.

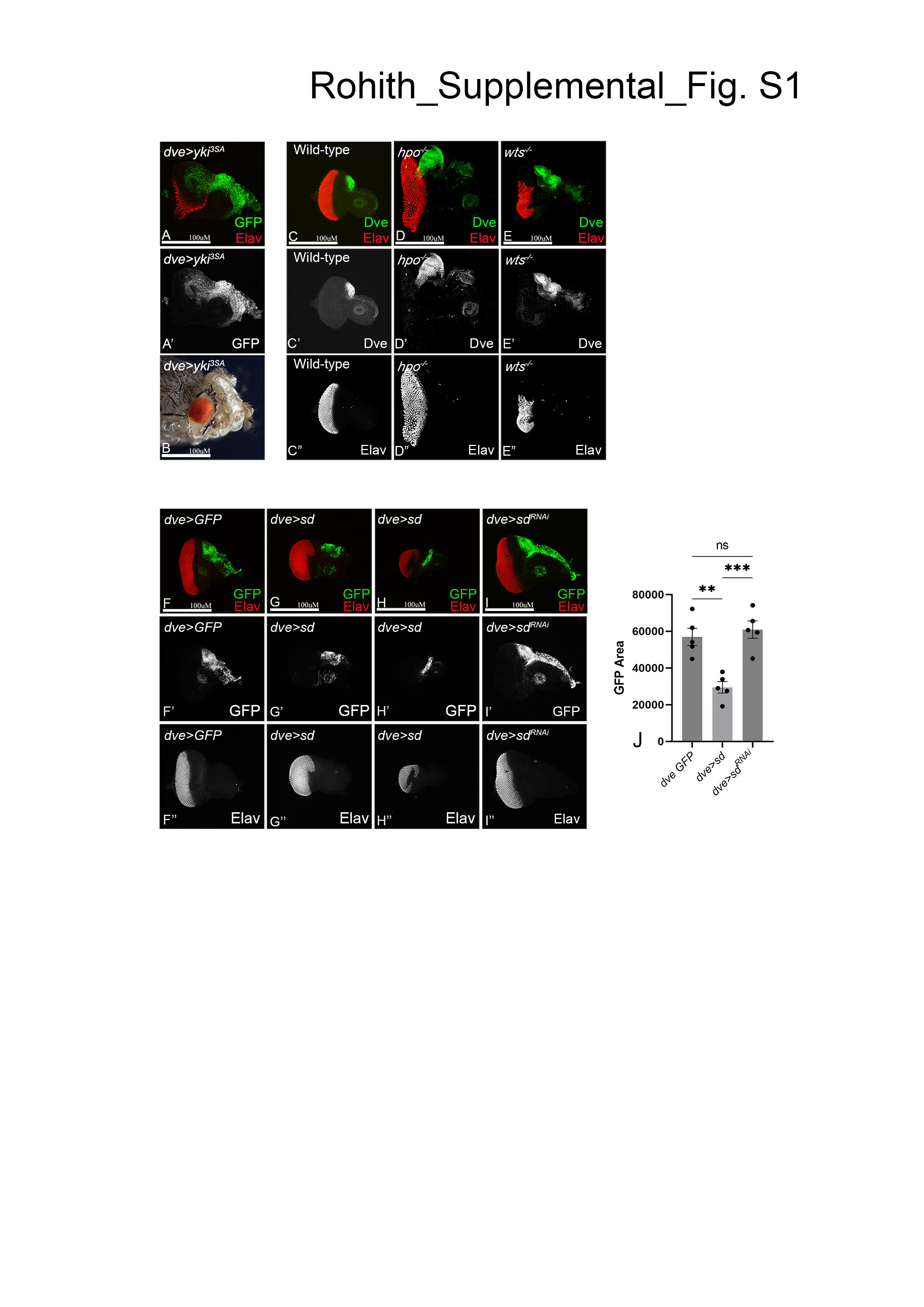

### Supplementary Figure S2 (linked to Figure 3): Dve modulation and its effects on Yki activity.

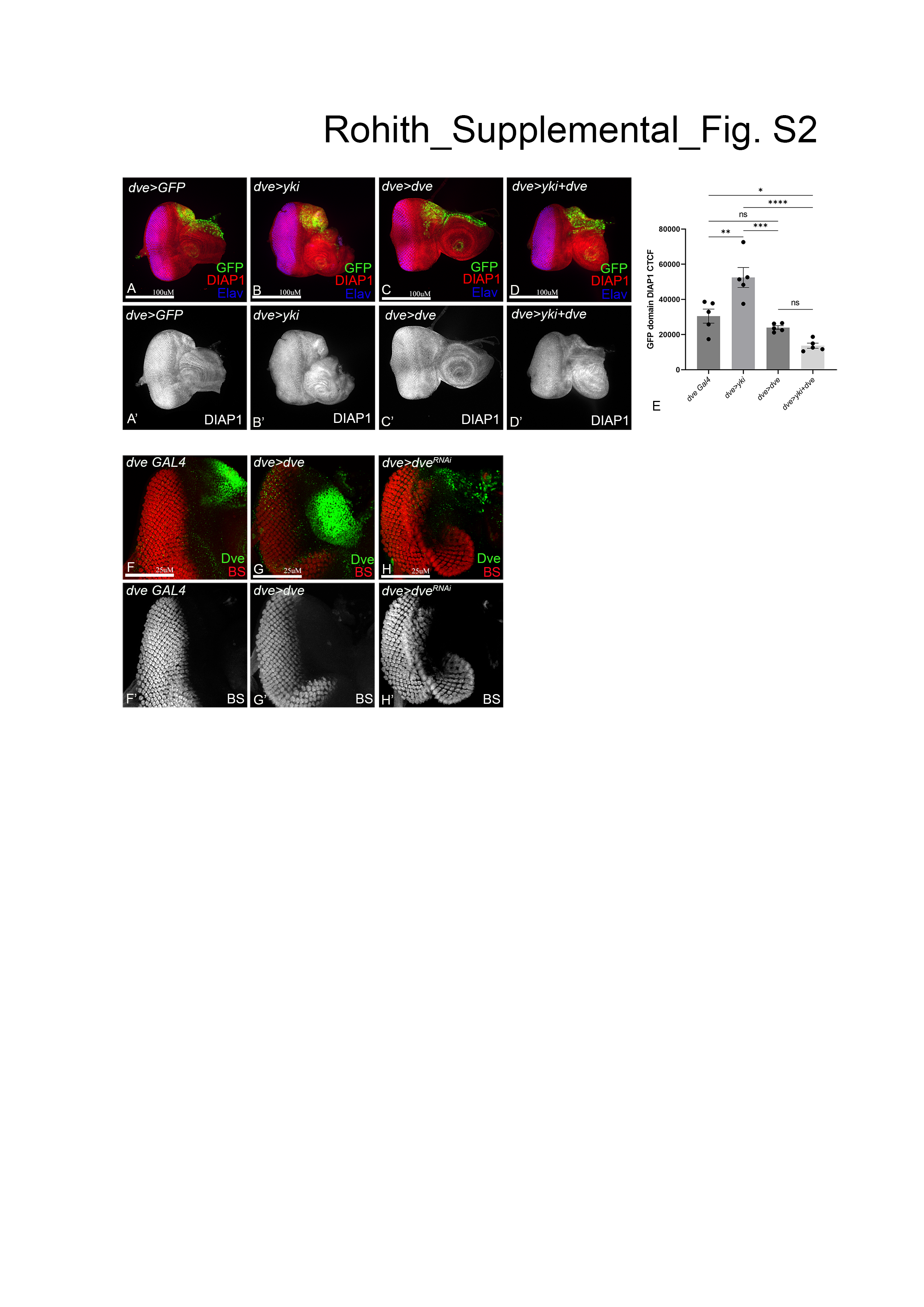

### Supplementary Figure S3 (linked to Figure 4): Yki does not transcriptionally regulate dve.

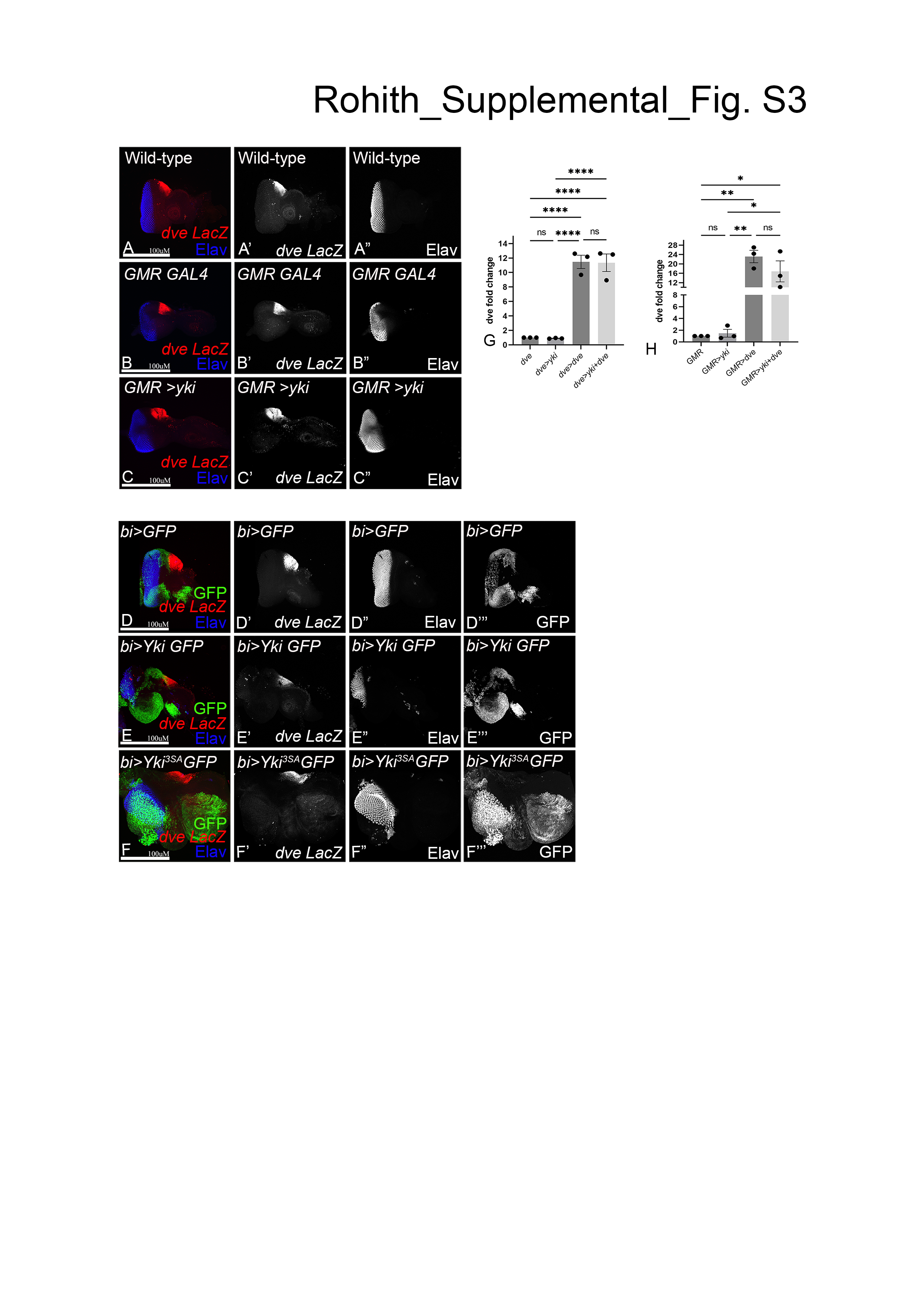

### Supplementary Figure S4 (linked to Figure 5): Effects on eye disc area and Wg expression in genetic epistasis interactions.

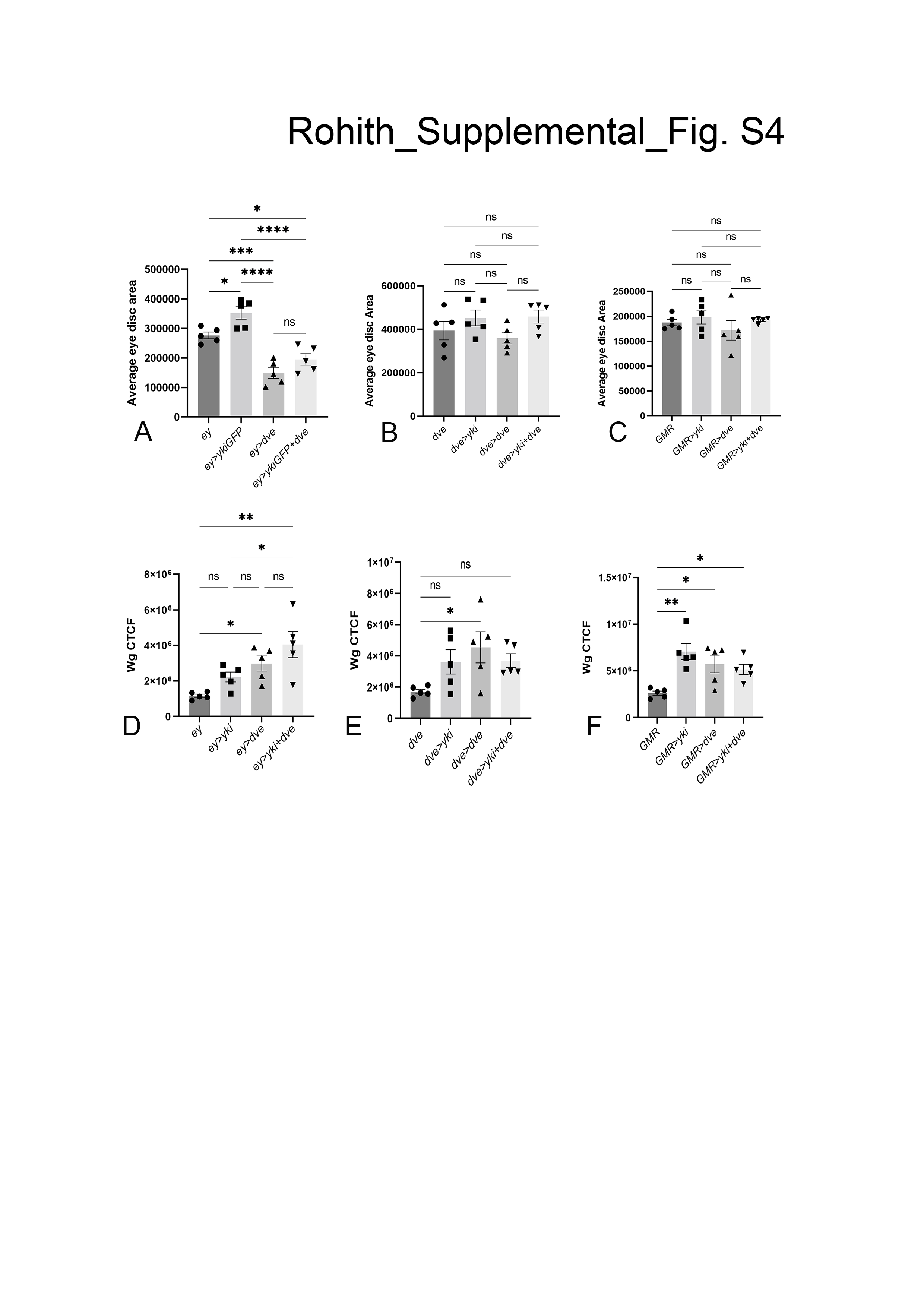
